# Blurred molecular epidemiological lines between the two dominant methicillin-resistant *Staphylococcus aureus* clones

**DOI:** 10.1101/501833

**Authors:** Amy C. Dupper, Mitchell J. Sullivan, Kieran I. Chacko, Aaron Mishkin, Brianne Ciferri, Ajay Kumaresh, Ana Berbel Caban, Irina Oussenko, Colleen Beckford, Nathalie E. Zeitouni, Robert Sebra, Camille Hamula, Melissa Smith, Andrew Kasarskis, Gopi Patel, Russell B. McBride, Harm van Bakel, Deena R. Altman

**Author notes:** Current affiliation is National Biodefense Analysis and Countermeasures Center (NBACC), 8300 Research Plaza, Fort Detrick, MD 21702. Address correspondence to: Deena Altman, MD, (corresponding author) or Amy Dupper, MPH, (alternate corresponding author).

## Abstract

**Background:** Methicillin-resistant *Staphylococcus aureus* (MRSA) causes life-threatening infections in both community and hospital settings and is a leading cause of healthcare-associated infections (HAIs). We sought to describe the molecular epidemiological landscape of patients with MRSA bloodstream infections (BSIs) at an urban medical center by evaluating the clinical characteristics associated with the two dominant endemic clones.

**Methods:** Comprehensive clinical data extraction from the electronic health records of 227 hospitalized patients ≥18 years old with MRSA BSI over a 33-month period in New York City were collected. The descriptive epidemiology and mortality associated with the two dominant clones was compared using logistic regression.

**Results:** Molecular analysis revealed that 91% of all single-patient MRSA BSIs were due to two equally represented genotypes, clonal complex (CC) 5 (N=117) and CC8 (N=110). MRSA BSIs were associated with a 90-day mortality of 27%. CC8 caused disease more frequently in younger age groups (56 ± 17 vs 67 ± 17 years old; *p*<0.001) and in non-White race (OR=3.45 95% CI [1.51-7.87]; p=0.003), with few other major distinguishing features. Morbidity and mortality also did not differ significantly between the two clones. CC8 caused BSIs more frequently in the setting of peripheral intravenous catheters (OR=5.96; 95% CI [1.51-23.50]; p=0.01).

**Conclusion:** The clinical features distinguishing dominant MRSA clones continue to converge. The association of CC8 with peripheral intravenous catheter infections underscores the importance of classical community clones causing hospital-onset infections. Ongoing monitoring and analysis of the dynamic epidemiology of this endemic pathogen is crucial to inform management to prevent disease.

## Introduction

Healthcare-associated infections (HAIs) pose a potentially fatal threat to patients worldwide^1^ and *Staphylococcus aureus* is one of the most common causes of HAIs in the United States.^2,3^ Methicillin-resistant *S. aureus* (MRSA) bloodstream infections (BSIs) are linked with mortality up to 30% and are associated with longer hospital stays and increased healthcare costs.^4,5^ MRSA has long been present in healthcare settings but is now well established in the community.^6^ The two dominant MRSA clones in the United States are clonal complex (CC) 5 and CC8.^3^ Historically, CC5 has been associated with older individuals with hospital or long-term care facility contact.^6^ In contrast, CC8, predominantly the USA300 pulsotype, was first reported in the US in healthy children in 1990s and raised concern for its capacity to cause severe disease in healthy individuals.^7^ Over the following two decades, CC8, largely driven by the USA300 lineage, became established as the predominant community-associated clone, presenting as skin and soft tissue infections (SSTIs) in athletes, children in day-care centers, injection drug users, and in persons with human immunodeficiency virus (HIV) infection.^8,9^

The prevalence of CC8 has increased in healthcare settings and is now associated with as many inpatient infections as CC5.^6,9^ In this connection, we sought to update and expand on the clinical aspects of the molecular epidemiology of MRSA BSIs in a major academic medical center in New York City. We examined the differences between the two dominant clonal complexes, CC5 and CC8, and their associated clinical and epidemiological features. We additionally studied clonal associations in the context of current surveillance definitions as defined by the National Healthcare Safety Network (NHSN), which are reportable.^10^ We explored clones in the context of associations with inpatient and outpatient community healthcare networks. Finally, we examined subgroups within the two CCs with an interest in the clinical features of the USA500, a relatively understudied clone.^11,12^ With extensive clinical detail we describe a picture more complex than genotypic associations are able to describe.

## Methods

### Study setting, patient identification, and molecular typing

The Mount Sinai Hospital (MSH) is a 1,018-bed tertiary- and quaternary-care facility. Under the approval of the MSH institutional review board, data were captured on a total of 249 adult (≥ 18 years old) patients with MRSA BSIs by the MSH Clinical Microbiology Laboratory as part of standard clinical care between August 2014 and April 2017. Identification and susceptibility of MRSA was performed using VITEK^®^2 (bioMerieux). From a hospital-wide genomic surveillance program, we derived the CC and multilocus sequence typing (MLST) *in silico* using the RESTful interface to the *S. aureus* PubMLST^13^ database. Staphylococcal protein A (*spa)* and Panton-Valentine Leukocidin (PVL) were generated using a custom script (https://github.com/mjsull/spa_typing) and BLAST+^14^ respectively. Core-genome MLST types were determined using the schema available at (https://www.cgmlst.org/ncs/schema/141106/). A tree for the visualization of clusters was created using GrapeTree,^15^ using representative published NCBI references (USA500: CP007499.1; USA100: GCA_000525105; USA300: NC_007793.1). The raw sequence data have been deposited in the National Center for Biotechnology Information SRA database under the Bioproject PRJNA470993.

### Patient data collection

Demographic and clinical data were obtained retrospectively from the electronic medical record system, including geographic admission data, presumed source of the BSI based on Infectious Diseases (ID) consultant, comorbidities, and prior outpatient healthcare exposures. All patients diagnosed with MRSA BSI received a consultation from an ID specialist at the time of diagnosis as per standard practice at our institution. The online database REDCap^16^ was used for data capture and to calculate the Charlson Comorbidity Index (CCI).^17^ Patients with non-CC5 or CC8 MRSA were excluded, resulting in a total of 227 patients for analyses. Zip codes were used to create a map of clones using the geographic information system (GIS) software ESRI Spatial Analysis.^18^ Surveillance definitions included hospital-onset MRSA (HO-MRSA), defined as positive cultures on or after the fourth day after hospital admission, and community-onset MRSA (CO-MRSA) defined as BSI presenting within the 72-hour hospital admission interval.^10^

### Statistical analysis

We selected established clinical correlates related to prior epidemiological studies, including demographics, baseline comorbidities, admission sources, and infection sources. We also evaluated in-hospital outcomes and death, especially those related to the MRSA BSI. Variables were collapsed to make the final set of covariates informative and reflective of current published literature. Analyses were performed in SAS (ver.9.4),^19^ and all figures were produced using R (ver.3.4.2).^20^ Non-normally distributed continuous variables were categorized into discrete categorical groups. Variables were first analyzed in a univariate logistic regression model, with variables *p*≤0.2 then placed into a multivariate logistic regression model. Mortality analyses were analyzed using a Cox regression model. All variables with *p*≤0.05 were considered statistically significant.

## Results

### Molecular composition of clones involved in MRSA BSI

Molecular analysis of single-patient, first episode MRSA BSI revealed the majority of MRSA BSIs were caused by the two dominant clones, CC5 and CC8 (Figure 1, Supplementary Table 1). CC8 was the cause of BSI in 110 (44%), and CC5 in 117 (47%) of the total of 249 cases, representing 91% of the entire population. Only 22 (9%) were non-CC5/CC8. Within CC5, the majority were either sequence type (ST) 5 (N=49; 42%) or ST105 (N=61; 52%). Six percent (N=7) belonged to other STs within CC5. The majority of CC8 isolates were ST8 (N=108; 98%) with 2 (2%) additional non-ST8 clones. The majority (N=64; 58%) of ST8 were *spa* type t008, along with 20 (18%) non-t008 *spa* types which also clustered with USA300. Additionally, CC8 included 24 (22%) *spa* type t064, including three (3%) non-t064 *spa* types, all clustering in the USA500 lineage.^11,21^

**Figure 1.**
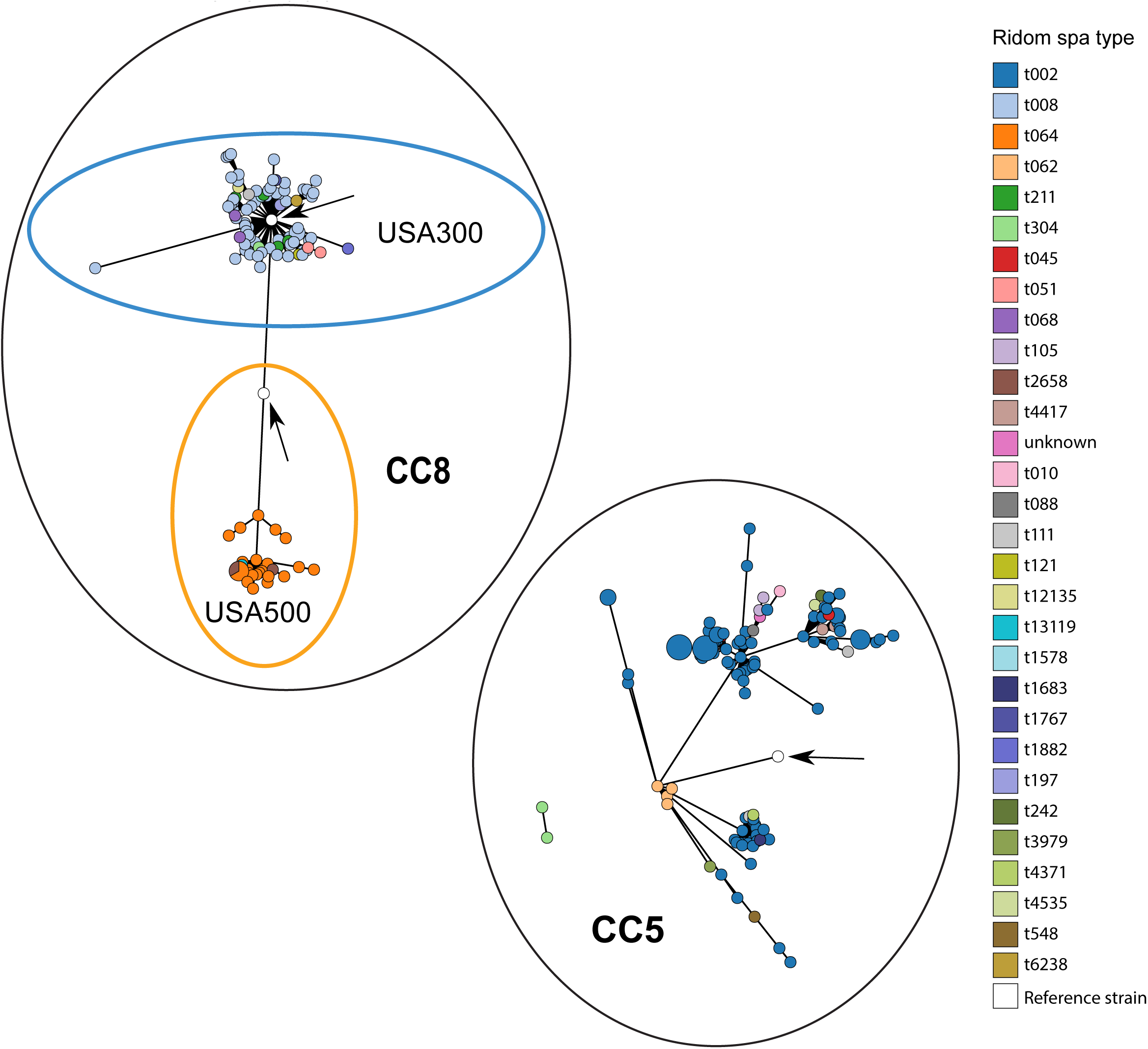
Core Genome Multilocus Sequence Typing (CG-MLST) of MRSA BSI isolates in study. Clustering of isolates into CC5 and CC8, as well as USA300 and USA500 based on cgMLST data. Ridom *spa* types are listed on <u>the</u> right. The arrows point to the published NCBI reference genomes for each grouping. “Unknown” refers to a *spa* type not listed in Ridom database.

**Figure 2.**
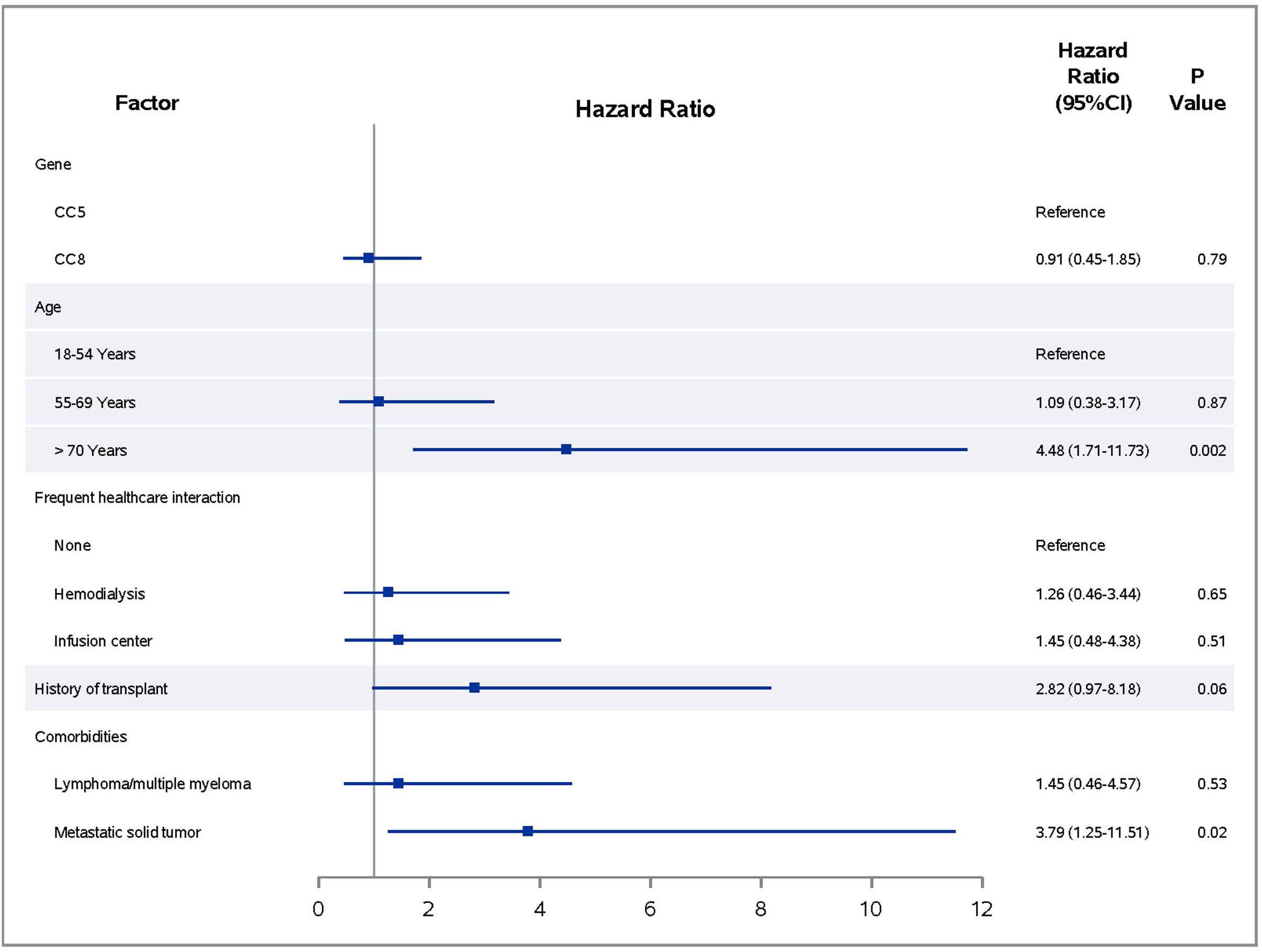
Multivariate analysis of death related to MRSA BSI within 90 days

### Baseline clinical characteristics of patients with MRSA BSIs

Sixty-seven percent of patients were male, and the median age at diagnosis was 62 years old (Table 1). The racial and ethnic composition included non-Hispanic White (N=98; 43%), non-Hispanic Black (N=63; 28%), Hispanic/Latino (N=46; 20%), Asian (N=8; 4%), and not reported (N=12; 5%). MRSA BSIs were linked to a wide range of causes, with vascular access (N=78; 34%), pneumonia (N=24; 11%), and SSTIs (N=25; 11%) representing the most common causes.

**Table 1.** Demographics and clinical characteristics of patients with MRSA BSIs

| Factor, n (%) | MRSA BSI<br>(n = 227) | Factor, n (%) | MRSA BSI<br>(n = 227) |
| --- | --- | --- | --- |
| *Clonal Complex* |  | *Comorbidities Continued* |  |
| CC8 | 110 (48) | Moderate/Severe Liver Disease | 17 (7) |
| CC5 | 117 (52) | Metastatic Solid Tumor | 14 (6) |
| *Sex* |  | *Charlson Comorbidity Index (CCI)* |  |
| Male | 151 (67) | 0-3 | 71 (31) |
| Female | 76 (33) | 4-5 | 48 (21) |
| *Race/Ethnicity* |  | 6-8 | 66 (29) |
| Non-Hispanic White | 98 (43) | > 8 | 42 (19) |
| Non-Hispanic Black | 63 (28) | <sup>g</sup>History of Transplant | 32 (14) |
| Hispanic/Latino | 46 (20) | <sup>h</sup>History of MRSA Colonization | 94 (41) |
| Asian | 8 (4) | *Presumed Source of MRSA BSI* |  |
| Unknown | 12 (5) | Peripheral Intravenous Catheter | 16 (7) |
| *Age at Time of Infection* |  | Skin & Soft Tissue Infection | 25 (11) |
| 18-54 Years | 79 (35) | Pneumonia | 24 (11) |
| 55-69 Years | 68 (30) | Diabetic Foot Infection | 17 (7) |
| ≥ 70 Years | 80 (35) | Vascular Access | 78 (34) |
| History of Injection Drug Use | 24 (11) | Septic Arthritis | 4 (2) |
| HIV | 22 (10) | Urinary Source | 4 (2) |
| *Admission Source* |  | Sacral Wound | 11 (5) |
| <sup>a</sup>Home | 132 (58) | Other/Unknown | 28 (12) |
| NH/Rehab/LTACH | 60 (26) | ICU Admission Prior to BSI | 42 (19) |
| Outside Hospital | 35 (15) |  |  |
| *Prior Hospital Admission (90 Days)* |  |  |  |
| *Length of Hospital Stay Prior to BSI* |  |  |  |
| ≤ 3 Days | 132 (58) |  |  |
| > 3 Days | 95 (42) |  |  |
| *Frequent Healthcare Interaction* |  |  |  |
| Hemodialysis | 40 (18) |  |  |
| <sup>b</sup>Infusion Center | 34 (15) |  |  |
| None | 153 (67) |  |  |
| <sup>c</sup>Presence of Invasive Device | 180 (79) |  |  |
| <sup>d</sup>Invasive Procedures | 109 (48) |  |  |
| <sup>e</sup>Wound Present | 94 (41) |  |  |
| <sup>f</sup>*Comorbidities* |  |  |  |
| Myocardial Infarction | 24 (11) |  |  |
| Congestive Heart Failure | 55 (24) |  |  |
| Peripheral Vascular Disease | 37 (16) |  |  |
| Cerebrovascular Disease | 20 (9) |  |  |
| Dementia | 29 (13) |  |  |
| Chronic Pulmonary Disease | 30 (13) |  |  |
| Connective Tissue Disease | 5 (2) |  |  |
| Peptic Ulcer Disease | 9 (4) |  |  |
| Mild Liver Disease | 2 (1) |  |  |
| Diabetes (no complications) | 38 (17) |  |  |
| Diabetes with Organ Damage | 51 (22) |  |  |
| Para or Hemiplegia | 17 (7) |  |  |
| Moderate/Severe Renal Disease | 55 (24) |  |  |
| Solid Tumor | 23 (10) |  |  |
| Leukemia | 13 (6) |  |  |

Abbreviations: LTACH, long-term acute care hospital; HIV, human immunodeficiency virus; BSI, blood stream infection; ICU, intensive care unit
<sup>a</sup> "Admission from home": included non-medical residences such as home, group homes, assisted living facilities, and homeless shelters.
<sup>b</sup> "Infusion center" : outpatient centers for chemotherapy, intravenous fluids, intravenous immunomodulators, and blood products.
<sup>c</sup> "Presence of an invasive device": included pacemaker, implantable cardioverter defibrillator (ICD), left ventricular assist device (LVAD), vascular access (excluding peripheral intravenous catheters), orthopedic hardware, nephrostomy, suprapubic catheter, ileal conduit, foley catheter, arteriovenous graft placement (AVG), percutaneous endoscopic gastrostomy (PEG) tube, or ostomy.
<sup>d</sup> "Invasive procedures": Included any invasive procedures or surgery occurring within one month prior to first positive blood culture for MRSA, excluding electroencephalogram (EEG), electrocardiogram (EKG), and transthoracic echocardiogram (TTE).
<sup>e</sup> "Wound Present": Presence of a chronic skin wound overlying the sacrum, limb, abdomen, or other body part.
<sup>f</sup> "Comorbidities": As defined by the Charleston Comorbidity Index (CCI) refer to standard definitions for CCI.<sup>17</sup>
<sup>g</sup> "History of transplant": Included solid organ and bone marrow transplant.
<sup>h</sup> "History of MRSA colonization": Any positive culture from urine, sputum, tissue, or nares with MRSA prior to the positive MRSA blood culture or a documented history of prior MRSA infection or colonization.

We performed a comprehensive analysis of admission sources. More than half of patients were admitted from home (N=128; 56%) with the remaining patients transferred in from nursing homes/rehabilitation/long-term care facilities (N=60; 26%), outside hospitals (N=35; 15%), or homeless shelters (N=4; 2%) (Table 1). Of subjects residing at home or group home settings, 32% (N=56) had frequent contact with healthcare centers either via outpatient dialysis (N=22; 12%) or infusion centers (N=34; 19%). Only 27 (12%) study patients had no significant inpatient or outpatient healthcare exposure.

The mean CCI of subjects on hospital admission was 5.4. The most common comorbid medical conditions in our dataset were congestive heart failure (N=55; 24%) and chronic renal disease (N=55; 24%). Ten percent (N=22) of patients were co-infected with HIV. Additionally, 32 (14%) had a history of a transplant (solid organ or bone marrow). Injection drug use was reported by 11% (N=24) of our population, and 41% (N=94) had a prior history of MRSA colonization.

### Clinical features and geographic distribution of the two dominant MRSA clones

As CC5 and CC8 were responsible for the majority of BSIs, we anchored our analyses on comparing these two clones. The majority of variables examined were not significantly increased in one clone over the other, with several notable exceptions. Race was found to confound the effects of HIV and injection drug use, so these two variables were retained in the final model. Logistic regression revealed that non-Hispanic Black race (OR=3.45; 95% CI [1.51-7.87]; *p*=0.003), Hispanic/Latino race (OR=3.21 95% CI [1.33-7.77]; *p*=0.01), HIV (OR=3.62 95% CI [1.01-12.92]; *p*=0.05), and SSTIs (OR=2.21; 95% CI [0.98-10.46]; *p*=0.05) had a higher likelihood of being infected with the CC8 (Table 2). Interestingly, CC8 was also increased in patients with peripheral intravenous catheters (PIVs) as the presumed source of MRSA BSI compared to CC5 (OR=5.96 95% CI [1.51-23.50]; *p*=0.01). Alternately, patients were less likely to have BSI with CC8 if they were greater than 70 years (OR=0.28 95% CI [0.11-0.74]; *p*=0.01) or if admitted from an outside hospital (OR=0.33 95% CI [0.11-1.00]; *p*=0.05) vs. home.

**Table 2.** Demographics and clinical characteristics of patients with MRSA BSIs stratified by clonal complex with the odds of being CC8 vs CC5

| Factor | CC8<br>N = 110 (%) | CC5<br>N = 117 (%) | Univariate Analysis |  | Multivariate Analysis |  |
| --- | --- | --- | --- | --- | --- | --- |
|  |  |  | OR (95% CI) | p value | OR (95% CI) | p value |
| Male | 73 (66) | 78 (67) | 0.99 (0.57-1.71) | 0.96 |  |  |
| *Race/Ethnicity* |  |  |  |  |  |  |
| Non-Hispanic White | 31 (28) | 67 (57) | Reference |  | Reference |  |
| Non-Hispanic Black | 41 (37) | 22 (19) | **4.03 (2.06-7.87)** | **<0.001** | **3.45 (1.51-7.87)** | **0.003** |
| Hispanic/Latino | 27 (25) | 19 (16) | **3.07 (1.49-6.34)** | **0.002** | **3.21(1.33-7.77)** | **0.01** |
| Asian | 4 (4) | 4 (3) | 2.16 (0.51-9.21) | 0.30 | 3.17 (0.61-16.38) | 0.17 |
| Unknown | 7 (6) | 5 (4) | 3.03 (0.89-10.29) | 0.08 | 2.14 (0.50-9.13) | 0.30 |
| *Age at Time of Infection* |  |  |  |  |  |  |
| 18-54 Years | 50 (45) | 29 (25) | Reference |  | Reference |  |
| 55-69 Years | 37 (34) | 31 (27) | 0.69 (0.36-1.34) | 0.28 | 0.64 (0.27-1.50) | 0.30 |
| ≥ 70 Years | 23 (21) | 57 (49) | **0.23 (0.12-0.46)** | **<0.001** | **0.28 (0.11-0.74)** | **0.01** |
| History of Injection Drug Use | 16 (15) | 8 (6) | 2.32 (0.95-5.66) | 0.06 | 0.62 (0.21-1.88) | 0.40 |
| HIV | 18 (16) | 4 (3) | **5.53 (1.81-16.90)** | **0.003** | **3.62 (1.01-12.92)** | **0.05** |
| *Admission Source* |  |  |  |  |  |  |
| Home | 78 (71) | 54 (46) | Reference |  | Reference |  |
| NH/Rehab/LTACH | 23 (21) | 37 (32) | **0.43 (0.23-0.80)** | **0.008** | 0.55 (0.24-1.26) | 0.16 |
| Other Hospital | 9 (8) | 26 (22) | **0.24 (0.10-0.55)** | **<0.001** | **0.33 (0.11-1.00)** | **0.05** |
| Prior Hospital Admission (90 Days) | 70 (64) | 92 (79) | **0.48 (0.26-0.86)** | **0.01** | 1.00 (0.43-2.32) | 0.99 |
| *Length of Hospital Stay Prior to BSI* |  |  |  |  |  |  |
| ≤3 Days | 74 (67) | 58 (50) | **2.09 (1.22-3.58)** | **0.007** | 2.01 (0.90-4.50) | 0.09 |
| >3 Days | 36 (33) | 59 (50) | Reference |  | Reference |  |
| *Frequent Healthcare Interaction* |  |  |  |  |  |  |
| Hemodialysis | 22 (20) | 18 (15) | 1.45 (0.72-2.92) | 0.30 |  |  |
| Infusion Center | 18 (16) | 16 (14) | 1.33 (0.63-2.81) | 0.45 |  |  |
| None | 70 (64) | 83 (71) | Reference |  |  |  |
| Presence of Invasive Device | 77 (70) | 103 (88) | **0.32 (0.16-0.63)** | **0.001** | 0.56 (0.24-1.32) | 0.18 |
| Invasive Procedures | 44 (40) | 65 (56) | **0.53 (0.32-0.90)** | **0.02** | 0.67 (0.30-1.48) | 0.32 |
| Wound Present | 46 (42) | 48 (41) | 1.03 (0.61-1.75) | 0.90 |  |  |
| *Charlson Comorbidity Index (CCI)* |  |  |  |  |  |  |
| 0-3 | 40 (36) | 31 (27) | Reference |  | Reference |  |
| 4-5 | 26 (24) | 22 (19) | 0.92 (0.44-1.91) | 0.82 | 1.34 (0.50-3.57) | 0.56 |
| 6-8 | 30 (27) | 36 (31) | 0.65 (0.33-1.27) | 0.20 | 1.09 (0.42-2.85) | 0.85 |
| >8 | 14 (13) | 28 (24) | **0.39 (0.18-0.86)** | **0.02** | 0.74 (0.22-2.49) | 0.63 |
| History of Transplant | 14 (13) | 18 (15) | 0.80 (0.38-1.70) | 0.57 |  |  |
| History of MRSA Colonization | 47 (43) | 47 (40) | 1.11 (0.66-1.89) | 0.70 |  |  |
| *Presumed Source of MRSA BSI* |  |  |  |  |  |  |
| Peripheral Intravenous Catheter | 11 (10) | 5 (4) | 2.49 (0.84-7.41) | 0.10 | **5.96 (1.51-23.50)** | **0.01** |
| Skin & Soft Tissue Infection | 18 (16) | 7 (6) | **3.08 (1.23-7.68)** | **0.02** | **2.21 (0.98-10.46)** | **0.05** |
| Pneumonia | 12 (11) | 12 (10) | 1.07 (0.46-2.50) | 0.87 |  |  |
| Diabetic Foot Infection | 8 (7) | 9 (8) | 0.94 (0.35-2.53) | 0.90 |  |  |
| Vascular Access | 35 (32) | 43 (37) | 0.80 (0.46-1.39) | 0.43 |  |  |
| Septic Arthritis | 1 (1) | 3 (3) | 0.35 (0.04-3.40) | 0.36 |  |  |
| Urinary Source | 1 (1) | 3 (3) | 0.35 (0.04-3.40) | 0.36 |  |  |
| Sacral Wound | 6 (5) | 5 (4) | 1.29 (0.38-4.36) | 0.68 |  |  |
| Other/Unknown | 12 (11) | 16 (14) | 0.77 (0.35-1.72) | 0.53 |  |  |
| ICU Admission Prior to BSI | 16 (15) | 26 (22) | 0.60 (0.30-1.18) | 0.14 | 1.47 (0.53-4.07) | 0.46 |

**Bold** = significant at ≤ 0.05
Abbreviations: NH, nursing home; rehab, rehabilitation facility; LTACH, long-term acute care hospital; BSI, bloodstream infection; HIV, human immunodeficiency virus; ICU, intensive care unit. See Table 1 for definitions.

On multivariate analysis, there were equal proportions of CC8 vs CC5 across patient admission sources. We further mapped the clones according to patient zip code and found no significant clustering aside from the area surrounding the hospital (Supplementary Figure 1).

### Clinical characteristics of patients with MRSA BSI due to clonal subgroups

Although we performed our top level analysis at the CC level, we additionally evaluated potential clinical differences between subgroups within the CCs. In CC8, a total of 27 CC8 isolates clustered with USA500, which is considered a healthcare-associated clone.^11,12,21^ We thus compared USA500 to USA300 and found overall few differences aside from higher proportion of USA500 in those with HIV (OR=6.61 95% CI [1.38-31.67]; *p*=0.02) (Supplementary Table 2). We also compared ST105 and ST5 lineages within CC5, which had few clinical distinctions (Supplementary Table 3). Stepwise removal of any of these subgroups did not impact results of our top level CC8 vs. CC5 analyses, thus they were retained in the analyses.

### Addressing surveillance definitions in endemic settings

Given the importance placed on reporting BSIs based on NHSN definitions, we sought to provide clinical detail to the BSIs in the context of these definitions.^10^ Overall there were more CO-MRSA BSIs (N=132; 58%) than HO-MRSA BSIs (N=95; 42%). Logistic regression revealed that patients characterized as CO-MRSA were more likely to have CC8 (OR=2.33 95% CI [1.03-5.27]; *p*=0.04), and more likely to be receiving hemodialysis at the time of infection (OR=4.62 95% CI [1.23-17.29]; *p*=0.02) (Table 3). Conversely, CO-MRSA was less likely to be associated with prior invasive procedures (OR=0.18 95% CI [0.08-0.40]; *p*=<0.001), MRSA BSI from a PIV (OR=0.11 95% CI [0.03-0.46]; *p*=0.002), MRSA BSI from vascular access (OR=0.36 95% CI [0.13-0.95]; *p*=0.04), and intensive care unit (ICU) admission prior to BSI (OR=0.13 95% CI [0.04-0.43]; *p*=<0.001). Only 27 (12%) patients had no clear healthcare exposure, thus 88% of our total patient population had healthcare risk factors prior to their MRSA BSI.

**Table 3.** Demographics and clinical characteristics of patients with MRSA BSIs stratified by NHSN definitions with the odds of having CO-MRSA vs HO-MRSA

| Factor | CO-MRSA<br>N = 132 (%) | HO-MRSA<br>N = 95 (%) | Univariate Analysis |  | Multivariate Analysis |  |
| --- | --- | --- | --- | --- | --- | --- |
|  |  |  | OR (95% CI) | p value | OR (95% CI) | p value |
| CC8 | 74 (56) | 36 (38) | **2.09 (1.22-3.58)** | **0.007** | **2.33 (1.03-5.27)** | **0.04** |
| Male | 90 (68) | 61 (64) | 1.19 (0.68-2.08) | 0.53 |  |  |
| *Race/Ethnicity* |  |  |  |  |  |  |
| Non-Hispanic White | 50 (38) | 48 (51) | Reference |  | Reference |  |
| Non-Hispanic Black | 40 (30) | 23 (24) | 1.67 (0.87-3.19) | 0.12 | 1.14 (0.44-3.00) | 0.79 |
| Hispanic/Latino | 30 (23) | 16 (17) | 1.80 (0.87-3.72) | 0.11 | 1.12 (0.36-3.43) | 0.85 |
| Asian | 6 (5) | 2 (2) | 2.88 (0.55-14.98) | 0.21 | 2.67 (0.36-19.62) | 0.33 |
| Unknown | 6 (5) | 6 (6) | 0.96 (0.29-3.18) | 0.95 | 1.08 (0.21-5.47) | 0.93 |
| *Age at Time of Infection* |  |  |  |  |  |  |
| 18-54 Years | 43 (33) | 36 (38) | Reference |  |  |  |
| 55-69 Years | 44 (33) | 24 (25) | 1.54 (0.79-2.99) | 0.21 |  |  |
| ≥ 70 Years | 45 (34) | 35 (37) | 1.08 (0.58-2.01) | 0.82 |  |  |
| History of Injection Drug Use | 18 (14) | 6 (6) | 2.34 (0.89-6.15) | 0.08 | 0.56 (0.15-2.14) | 0.40 |
| HIV | 13 (10) | 9 (9) | 1.04 (0.43-2.55) | 0.93 |  |  |
| *Admission Source* |  |  |  |  |  |  |
| Home | 80 (61) | 52 (55) | Reference |  | Reference |  |
| NH/Rehab/LTACH | 40 (30) | 20 (21) | 1.30 (0.69-2.47) | 0.42 | 0.89 (0.34-2.30) | 0.81 |
| Other Hospital | 12 (9) | 23 (24) | **0.34 (0.16-0.74)** | **0.007** | 0.78 (0.25-2.46) | 0.67 |
| Prior Hospital Admission (90 Days) | 91 (69) | 71 (75) | 0.75 (0.42-1.36) | 0.34 |  |  |
| *Frequent Healthcare Interaction* |  |  |  |  |  |  |
| Hemodialysis | 33 (25) | 7 (7) | **3.48 (1.45-8.36)** | **0.005** | **4.62 (1.23-17.29)** | **0.02** |
| Infusion Center | 11 (8) | 23 (24) | **0.35 (0.16-0.78)** | **0.01** | 0.35 (0.12-1.07) | 0.07 |
| None | 88 (67) | 65 (68) | Reference |  | Reference |  |
| Presence of Invasive Device | 95 (72) | 85 (89) | **0.30 (0.14-0.64)** | **0.002** | 0.54 (0.19-1.55) | 0.25 |
| Invasive Procedures | 37 (28) | 72 (76) | **0.12 (0.07-0.23)** | **<0.001** | **0.18 (0.08-0.40)** | **<0.001** |
| Wound Present | 58 (44) | 36 (38) | 1.29 (0.75-2.20) | 0.36 |  |  |
| *Charlson Comorbidity Index (CCI)* |  |  |  |  |  |  |
| 0-3 | 38 (29) | 33 (35) | Reference |  | Reference |  |
| 4-5 | 24 (18) | 24 (25) | 0.87 (0.42-1.81) | 0.71 | 1.48 (0.51-4.23) | 0.47 |
| 6-8 | 40 (30) | 26 (27) | 1.34 (0.68-2.64) | 0.40 | 1.08 (0.40-2.92) | 0.88 |
| >8 | 30 (23) | 12 (13) | 2.17 (0.96-4.91) | 0.06 | 2.13 (0.62-7.25) | 0.23 |
| History of Transplant | 13 (10) | 19 (20) | **0.44 (0.20-0.94)** | **0.03** | 1.11 (0.38-3.23) | 0.85 |
| History of MRSA Colonization | 58 (44) | 36 (38) | 1.29 (0.75-2.20) | 0.36 |  |  |
| *Presumed Source of MRSA Infection* |  |  |  |  |  |  |
| Peripheral Intravenous Catheter | 6 (5) | 10 (11) | 0.41 (0.14-1.16) | 0.09 | **0.11 (0.03-0.46)** | **0.002** |
| Skin & Soft Tissue Infection | 17 (13) | 8 (8) | 1.61 (0.66-3.90) | 0.29 |  |  |
| Pneumonia | 14 (11) | 10 (11) | 1.01 (0.43-2.38) | 0.98 |  |  |
| Diabetic Foot Infection | 15 (11) | 2 (2) | **5.96 (1.33-26.73)** | **0.02** | 2.04 (0.33-12.45) | 0.44 |
| Vascular Access | 36 (27) | 42 (44) | **0.47 (0.27-0.83)** | **0.009** | **0.36 (0.13-0.95)** | **0.04** |
| Septic Arthritis | 2 (2) | 2 (2) | 0.72 (0.10-5.17) | 0.74 |  |  |
| Urinary Source | 3 (2) | 1 (1) | 2.18 (0.22-21.30) | 0.50 |  |  |
| Sacral Wound | 9 (7) | 2 (2) | 3.40 (0.72-16.12) | 0.12 | 0.70 (0.11-4.55) | 0.71 |
| Other/Unknown | 16 (12) | 12 (13) | 0.95 (0.43-2.12) | 0.91 |  |  |
| ICU Admission Prior to BSI | 5 (4) | 37 (39) | **0.06 (0.02-0.17)** | **<0.001** | **0.13 (0.04-0.43)** | **<0.001** |

**Bold** = significant at $\leq 0.05$
Abbreviations: NH, nursing home; rehab, rehabilitation facility; LTACH, long-term acute care hospital; BSI, bloodstream infection; HIV, human immunodeficiency virus; ICU, intensive care unit; NHSN, National Healthcare Safety Network. See Table 1 for definitions.

We also examined CC8 vs CC5 in the context of the NHSN definitions. CC8 was responsible for 38% of HO-MRSA BSIs and was associated with younger age and non-Hispanic Black race (Supplementary Tables 4 & 5). Among patients grouped into the HO-MRSA stratum, those whose MRSA BSI resulted from a PIV (OR=9.84 95% CI [1.46-66.50]; *p*=0.02) were more likely to be from CC8. We also explored clones and definitions in the context of the individual comorbidities that constitute the CCI, and we found that those with lymphoma and/or multiple myeloma (OR=0.30 95% CI [0.11-0.80]; *p*=0.02) were more likely to occur among HO-MRSA, yet in the CO-MRSA stratum, lymphoma and/or multiple myeloma was more likely to involve CC5 (OR=0.05 95% CI [0.01-0.48]; *p*=0.01).

### Differences in morbidity and mortality related to MRSA BSIs

Morbidity outcomes associated with MRSA BSI such as need for ICU admission, need for mechanical ventilation, and development of metastatic complications were studied with respect to clone. Overall, we found no differences in morbidity between the two clones (Supplementary Table 6C & D). Interestingly, strictly CO-MRSA had overall worse hospital outcomes, with increased ICU admissions (OR=10.73 95% CI [3.94-29.26]; *p*=<0.001), mechanical ventilation (OR=3.45 95% CI [1.30-9.15]; *p*=0.01), and metastatic complications (OR=3.34 95% CI [1.46-7.64]; *p*=0.004) associated with MRSA BSI (Supplementary Table 6B). Overall, 20% (N=46) had persistent bacteremia (defined as BSI lasting >7 days) with no clonal predominance in these cases.

With regard to mortality, we examined 90-day all-cause and 90-day mortality associated with MRSA BSI. All-cause 90-day mortality was 27% (N=61) and of those that died at 90-days, death was associated with MRSA BSI in 54% (N=33) of cases. All-cause 90-day mortality had a lower likelihood of death due to CC8 vs. CC5 (OR=0.55 95% CI [0.30-1.00]; *p*=0.05), which was also observed in the CO-MRSA stratum (OR=0.43 95% CI [0.19-0.99]; *p*=0.05).

As a correlate for pathogenesis, we examined whether one clone had higher 90-day mortality in the setting of MRSA BSI. We first looked solely at the survival curves of each clone (Supplementary Figure 2), which revealed no difference. Second, we examined the clone variable in a multivariate Cox regression with possible confounders (Supplementary Figure 3). Interestingly, there was no difference in MRSA-related 90-day mortality related to MRSA with respect to clones (OR=0.91 95% CI [0.45-1.85]; *p*=0.79). Ninety-day mortality related to MRSA was more likely to occur in individuals older than 70 years (OR=4.48 95% CI [1.71-11.73]; *p*=0.002) and those with metastatic solid tumors (OR=3.79 95% CI [1.25-11.51]; *p*=0.02). Finally, we examined 90-day mortality associated with MRSA BSI due to primary sources of bacteremia, which revealed higher mortality with MRSA BSI from pneumonia (OR=2.93 95% CI [0.98-8.74]; *p*=0.05) or septic arthritis (OR=5.80 95% CI [1.05-32.13]; *p*=0.04).

## Discussion

As clones causing invasive MRSA infections are tied to specific populations, syndromes and settings, and are thought to behave differently, we sought to unravel how these associations manifest clinically in BSIs in a high level care institution in an endemic region. This study represents a large cohort of patients who were selected based strictly on presence of invasive disease (bacteremia) and demonstrate highly complex cases linked to significant morbidity and mortality. Consistent with previous reports from our region^22,23^ and across the United States,^3^ the majority of isolates were either CC5 or CC8. We demonstrated that CC8, representing half of all MRSA BSIs, was more frequently seen in younger age and non-White race. These data support the described convergence of clinical features classically associated with the two dominant clones.^6,24^ Although stratification by surveillance definitions was consistent with clones to an extent, it questions the applicability of definitions in endemic regions. These findings provide support for the concept that classic community genotypes involve individuals with frequent healthcare interactions.^25^

Although CC8/USA300 has been considered hypervirulent, by causing disease in younger, healthier individuals,^24,26,27^ work performed in animal models does not always actualize in complex human infections.^28,29^ Our study did not find significant differences in mortality and other outcomes based on MRSA clone, even after adjusting for comorbidities. This suggests that apparent differences driving morbidity and mortality are not solely due to differences between genotypes but due to a complex combination of demographic, host, and genomic factors, which require further study in human populations. Patients in our study had a CCI of 5.4, which is higher than most other studies that cite ranges of 1.5-3.^17,24^ We found increased mortality in the setting of MRSA BSI among those older than 70 years and those with metastatic solid tumors. Although these underlying conditions are independently associated with an increased risk for death, these data suggest that these populations have worse outcomes over other comorbid conditions when they develop MRSA BSI, and are hence a focus in future interventions. Finally, a higher proportion of males had MRSA BSI in our study, consistent with prior studies.^30,31^

Of interest was the increase in BSIs due to the USA500 in persons with HIV (PWH), a finding echoed in recent abstracts.^32,33^ This suggests that the types of MRSA infecting PWH may be shifting away from the historical USA300.^6^ As USA500 is considered a healthcare clone,^6,11,12^ this may reflect the changing epidemiology and management of HIV as it becomes a more chronic condition.

We describe increased CC8 in PIV-related BSIs, which was also significant in the HO-MRSA stratum. Source determination was based on the documentation of thrombophlebitis in all cases, as described in detail by ID consultants. We did not label the PIV as source unless it was clearly stated by ID clinicians to be the actual source, and ensured these infections were not incorrectly categorized as other types of skin infections. PIV placement is an aseptic but not a sterile procedure, and more emphasis and attention is placed on maintenance of central venous catheters than on PIVs. Although the incidence of PIV-related BSIs is low, the high frequency of PIV use results in a significant portion of PIVs resulting in BSIs.^34^ BSIs derive from colonizing flora,^35^ and the CC8 (USA300 in particular) is associated with skin colonization,^36^ which is supported in our data by the increase in CC8 in the setting of SSTIs. For the PIV infections and CC8 association in the HO-MRSA stratum, the most likely explanation is that patients are already colonized with CC8 and subsequently become infected with their isolate after PIV-related complications.^37^ This work further builds upon community origins of HO-MRSA BSIs by adding the association of CC8 with PIVs among patients with HO-MRSA.^38^ A larger sample size and access to colonizing isolates would assist in expansion of this concept, highlight under-recognized HAIs,^30^ and assist in evaluating the role of patient hygiene.

This study also examined the challenges of current surveillance definitions to describe these infections. A striking 88% of all patients had previous healthcare exposures, and 80% of the strictly CO-MRSA had prior healthcare exposures. An alternative definition of community-onset healthcare-associated (CO-HCA) for BSIs that present within three days of hospital admission in patients with frequent healthcare exposure^4,26^ may be more appropriate to report. We also describe that those presenting with BSIs from the community are a medically complex group with poor outcomes, consistent with studies associating CO-MRSA with complicated bacteremia.^39^ Furthermore, those infected with the CO-MRSA with CC5 were well advanced in their disease course, either through delay of presentation to the hospital or through transfers from other facilities for advanced care (as noted that 52 (39%) of admissions in the CO-MRSA group were admitted from other facilities). It appears that future descriptions of these classifications should include the changing epidemiology of the patients and their complex medical experiences.

This study has several limitations. As a retrospective study, it is subject to errors in chart abstraction. Being a single institution study, findings from the medically complex population studied may not be generalizable to smaller community hospitals. Our primary endpoint of mortality may be subject to reporting bias since death occurring outside the hospital may not have been captured in medical records. Although we recognize that methicillin-susceptible *S. aureus* (MSSA) also causes significant disease, we focused on MRSA-BSI due to the focus of our molecular surveillance program.^40^ Similar analyses extending beyond BSIs and including MSSA are critical.

In a highly complex patient population, there remain few distinct differences in the characteristics associated with the two endemic clones. CC8 has become even more common in the hospital, and it behaves similarly to CC5 by infecting infirm individuals. There were likewise no significant differences in both morbidity and mortality outcomes. Our study also highlights shifts in molecular epidemiology, at-risk populations, and potential areas for prevention such as PIVs in order to forestall this fatal disease. Integration of these clinical correlates with genomic and other multiscale analyses will lead to a more complete understanding *S. aureus* pathogenesis.

