## Supplementary Materials for "Blurred molecular epidemiological lines between the two dominant methicillin-resistant *Staphylococcus aureus* clones"

### SUPPLEMENTAL MATERIALS

#### Supplementary Table 1. Subject ID numbers, sample accession numbers and statistics, and basic clonal markers for MRSA isolates in study

| REDCap ID | Clonal Complex | MLST | Spa Type | PVL | SRR Accession | Read Number | Average Read Length |
|---|---|---|---|---|---|---|---|
| 1 | CC5 | 105 | t002 | -/- | SRR9722754 | 77658 | 6270.8 |
| 2 | CC5 | 105 | t002 | -/- | SRR9722862 | 143433 | 6664.1 |
| 3 | CC5 | 105 | t002 | -/- | SRR9722832 | 143773 | 6035.3 |
| 4 | CC5 | 105 | t002 | -/- | SRR9722864 | 161920 | 7134.1 |
| 5 | CC5 | 105 | t002 | -/- | SRR9722777 | 173197 | 7474.7 |
| 6 | CC5 | 105 | t002 | -/- | SRR9722873 | 141051 | 7310.7 |
| 7 | CC5 | 105 | t002 | -/- | SRR9722892 | 192647 | 7117.6 |
| 8 | CC5 | 105 | t002 | -/- | SRR9722891 | 199141 | 7200.0 |
| 9 | CC5 | 105 | t002 | -/- | SRR9722827 | 174234 | 6720.9 |
| 10 | CC5 | 105 | t002 | -/- | SRR9722831 | 173686 | 6408.5 |
| 11 | CC5 | 5 | t002 | -/- | SRR9722610 | 129297 | 7327.4 |
| 12 | CC8 | 8 | t008 | +/+ | SRR9722642 | 159161 | 6651.4 |
| 13 | CC5 | 105 | t002 | -/- | SRR9722595 | 138973 | 6802.7 |
| 14 | CC5 | 5 | t002 | -/- | SRR9722587 | 116526 | 5313.7 |
| 15 | CC5 | 105 | t002 | -/- | SRR9722631 | 145971 | 7295.9 |
| 16 | CC5 | 105 | t002 | -/- | SRR9722663 | 174366 | 7548.8 |
| 17 | CC5 | 105 | t002 | -/- | SRR9722671 | 153786 | 6038.3 |
| 18 | CC8 | 8 | t064 | -/- | SRR9722652 | 163843 | 6428.0 |
| 19 | CC5 | 105 | t002 | -/- | SRR9722585 | 156906 | 7454.9 |
| 20 | CC5 | 105 | t002 | -/- | SRR9722635 | 165663 | 7081.2 |
| 21 | CC8 | 8 | t2658 | -/- | SRR9722866 | 150161 | 7127.2 |
| 22 | CC8 | 8 | t064 | -/- | SRR9722860 | 170674 | 7419.7 |
| 23 | CC5 | 5 | t002 | -/- | SRR9722856 | 151277 | 6872.6 |
| 25 | CC8 | 8 | t008 | -/- | SRR9722603 | 118490 | 6640.3 |
| 26 | CC5 | 5 | t002 | -/- | SRR9722604 | 117166 | 5962.2 |
| 27 | CC8 | 8 | t008 | +/+ | SRR9722605 | 114230 | 4550.9 |
| 28 | CC5 | 105 | t002 | -/- | SRR9722606 | 126383 | 6468.8 |
| 29 | CC8 | 8 | t008 | +/+ | SRR9722599 | 110158 | 7506.4 |
| 30 | CC5 | 105 | t002 | -/- | SRR9722600 | 136094 | 6835.1 |
| 32 | CC5 | 105 | t002 | -/- | SRR9722601 | 103046 | 5833.2 |
| 33 | CC5 | 105 | t002 | -/- | SRR9722607 | 107199 | 6612.0 |
| 34 | CC5 | 5 | t002 | -/- | SRR9722548 | 101164 | 5890.3 |
| 35 | CC8 | 8 | t064 | -/- | SRR9722817 | 114987 | 5377.3 |
| 36 | CC5 | 105 | t002 | -/- | SRR9722550 | 119906 | 7121.5 |
| 37 | CC5 | 105 | t088 | -/- | SRR9722551 | 88986 | 4387.6 |
| 38 | CC5 | 105 | t002 | -/- | SRR9722555 | 108165 | 5389.3 |
| 39 | CC5 | 105 | t002 | -/- | SRR9722552 | 237799 | 5677.8 |
| 40 | CC8 | 8 | t008 | +/+ | SRR9722553 | 115197 | 7026.0 |
| 41 | CC8 | 8 | t008 | +/+ | SRR9722554 | 130521 | 7016.4 |
| 42 | CC5 | 5 | t002 | -/- | SRR9722556 | 139536 | 6992.6 |
| 43 | CC5 | 105 | t002 | -/- | SRR9722557 | 94231 | 4371.9 |
| 44 | CC5 | 5 | t002 | -/- | SRR9722575 | 154999 | 6931.4 |
| 45 | CC8 | 8 | t13119 | -/- | SRR9722574 | 34857 | 4578.7 |
| 47 | CC5 | 496 | t002 | -/- | SRR9722579 | 126044 | 6131.6 |

| REDCap ID | Clonal Complex | MLST | Spa Type | PVL | SRR Accession | Read Number | Average read length |
|---|---|---|---|---|---|---|---|
| 48 | CC5 | 496 | t002 | -/- | SRR9722570 | 98279 | 5797.5 |
| 49 | CC5 | 105 | t002 | -/- | SRR9722569 | 106487 | 5987.1 |
| 50 | CC8 | 8 | t008 | -/+ | SRR9722814 | 15865 | 4277.0 |
| 51 | CC5 | 105 | t002 | -/- | SRR9722573 | 95153 | 5690.2 |
| 52 | CC5 | 105 | t002 | -/- | SRR9722568 | 120471 | 5447.3 |
| 53 | CC8 | 8 | t008 | +/+ | SRR9722571 | 167518 | 6896.8 |
| 54 | CC8 | 8 | t304 | +/+ | SRR9722645 | 185301 | 6919.2 |
| 56 | CC8 | 8 | t008 | -/- | SRR9722643 | 163137 | 7015.8 |
| 57 | CC8 | 8 | t008 | +/+ | SRR9722641 | 166128 | 6788.6 |
| 58 | CC5 | 5 | t002 | -/- | SRR9722639 | 160300 | 6626.7 |
| 59 | CC8 | 8 | t008 | +/+ | SRR9722640 | 190344 | 6389.0 |
| 60 | CC5 | 5 | t062 | -/- | SRR9722637 | 172017 | 5625.8 |
| 61 | CC5 | 5 | t002 | -/- | SRR9722638 | 189600 | 7038.9 |
| 63 | CC8 | 8 | t121 | +/+ | SRR9722589 | 156387 | 5821.9 |
| 64 | CC8 | 8 | t008 | +/+ | SRR9722588 | 197575 | 5798.2 |
| 65 | CC8 | 8 | t008 | +/+ | SRR9722593 | 116277 | 4395.7 |
| 66 | CC8 | 8 | t064 | -/- | SRR9722592 | 144238 | 6914.3 |
| 67 | CC8 | 8 | t211 | +/+ | SRR9722591 | 156224 | 6589.6 |
| 68 | CC5 | 105 | t105 | -/- | SRR9722586 | 100986 | 4125.3 |
| 69 | CC8 | 8 | t211 | +/+ | SRR9722614 | 121587 | 5724.2 |
| 70 | CC8 | 8 | t008 | +/+ | SRR9722616 | 162610 | 6402.6 |
| 71 | CC8 | 8 | t068 | +/+ | SRR9722608 | 74052 | 4408.3 |
| 73 | CC8 | 8 | t008 | +/+ | SRR9722611 | 135636 | 6375.5 |
| 74 | CC5 | 105 | t002 | -/- | SRR9722612 | 85166 | 4015.4 |
| 75 | CC5 | 5 | t002 | -/- | SRR9722617 | 117779 | 6919.1 |
| 76 | CC8 | 8 | t008 | +/+ | SRR9722618 | 137775 | 6598.8 |
| 77 | CC8 | 8 | t008 | +/+ | SRR9722629 | 176878 | 6989.5 |
| 79 | CC8 | 8 | t008 | +/+ | SRR9722634 | 163260 | 5160.4 |
| 80 | CC8 | 8 | t008 | +/+ | SRR9722658 | 145455 | 6058.7 |
| 81 | CC5 | 5 | t062 | -/- | SRR9722657 | 149432 | 6445.0 |
| 83 | CC5 | 5 | t002 | -/- | SRR9722659 | 178910 | 7526.7 |
| 84 | CC5 | 105 | t002 | -/- | SRR9722664 | 179052 | 7640.9 |
| 85 | CC5 | 5 | t002 | -/- | SRR9722661 | 160348 | 7688.3 |
| 86 | CC5 | 5 | t002 | -/- | SRR9722662 | 154481 | 7731.8 |
| 87 | CC8 | 8 | t064 | -/- | SRR9722667 | 189141 | 7357.1 |
| 88 | CC5 | 225 | t668 | -/- | SRR9722668 | 191626 | 7115.7 |
| 89 | CC8 | 8 | t064 | -/- | SRR9722665 | 207540 | 6982.1 |
| 90 | CC8 | 8 | t008 | +/+ | SRR9722670 | 148647 | 6690.2 |
| 91 | CC5 | 105 | t002 | -/- | SRR9722546 | 191022 | 7550.5 |
| 92 | CC8 | 8 | t197 | +/+ | SRR9722545 | 184714 | 7598.6 |
| 94 | CC5 | 5 | t002 | -/- | SRR9722676 | 169633 | 6856.4 |
| 95 | CC8 | 8 | t008 | +/+ | SRR9722675 | 173401 | 6370.5 |
| 96 | CC5 | 105 | t4535 | -/- | SRR9722674 | 98750 | 6781.2 |
| 97 | CC8 | 8 | t008 | +/+ | SRR9722673 | 175297 | 6629.9 |
| 98 | CC8 | 8 | t008 | +/+ | SRR9722672 | 173480 | 6357.8 |
| 99 | CC8 | 8 | t12135 | +/+ | SRR9722560 | 209537 | 7094.5 |
| 100 | CC5 | 105 | t002 | -/- | SRR9722561 | 183590 | 7106.6 |

| REDCap ID | Clonal Complex | MLST | Spa Type | PVL | SRR Accession | Read Number | Average read length |
|---|---|---|---|---|---|---|---|
| 101 | CC8 | 8 | t008 | +/+ | SRR9722564 | 208585 | 7134.2 |
| 102 | CC5 | 105 | t002 | -/- | SRR9722565 | 201411 | 7277.1 |
| 103 | CC8 | 8 | t008 | +/+ | SRR9722566 | 162032 | 6746.2 |
| 104 | CC8 | 8 | t008 | +/+ | SRR9722567 | 211865 | 6791.0 |
| 105 | CC8 | 8 | t064 | -/- | SRR9722558 | 182009 | 6242.8 |
| 107 | CC5 | 5 | t002 | -/- | SRR9722654 | 255813 | 5872.1 |
| 108 | CC8 | 8 | t008 | +/+ | SRR9722653 | 204569 | 6697.8 |
| 109 | CC8 | 8 | t008 | -/+ | SRR9722651 | 213384 | 7023.1 |
| 110 | CC5 | 105 | t002 | -/- | SRR9722650 | 211273 | 6863.9 |
| 111 | CC8 | 8 | t008 | +/+ | SRR9722649 | 208857 | 6304.1 |
| 112 | CC8 | 8 | t064 | -/- | SRR9722647 | 182164 | 6720.1 |
| 113 | CC5 | 5 | t002 | +/+ | SRR9722580 | 169574 | 7025.7 |
| 114 | CC8 | 8 | t008 | +/+ | SRR9722581 | 150626 | 7021.5 |
| 115 | CC8 | 8 | t064 | -/- | SRR9722582 | 139316 | 6355.1 |
| 116 | CC5 | 105 | t002 | -/- | SRR9722583 | 160437 | 7282.6 |
| 117 | CC8 | 8 | t008 | +/+ | SRR9722636 | 159084 | 6595.0 |
| 119 | CC5 | 5 | t002 | -/- | SRR9722577 | 148894 | 7217.6 |
| 120 | CC5 | 5 | t242 | -/- | SRR9722578 | 170658 | 6948.5 |
| 121 | CC8 | 8 | t008 | +/- | SRR9722620 | 171104 | 6718.0 |
| 122 | CC5 | 5 | t002 | -/- | SRR9722619 | 172609 | 6674.3 |
| 123 | CC5 | 3390 | t002 | -/- | SRR9722622 | 136532 | 7367.3 |
| 124 | CC5 | 231 | t002 | -/- | SRR9722621 | 120997 | 7133.4 |
| 125 | CC8 | 8 | t008 | -/- | SRR9722624 | 156060 | 7353.1 |
| 127 | CC8 | 8 | t008 | +/+ | SRR9722626 | 119815 | 7324.8 |
| 128 | CC8 | 8 | t064 | -/- | SRR9722627 | 150085 | 7337.4 |
| 129 | CC8 | 8 | t064 | -/- | SRR9722597 | 133889 | 6884.3 |
| 130 | CC5 | 105 | t002 | -/- | SRR9722598 | 170321 | 7320.9 |
| 131 | CC8 | 8 | t008 | +/+ | SRR9722596 | 119227 | 7382.1 |
| 132 | CC8 | 8 | t008 | -/- | SRR9722882 | 166444 | 7127.0 |
| 134 | CC5 | 5 | t002 | -/- | SRR9722802 | 161990 | 7299.7 |
| 135 | CC5 | 105 | t002 | -/- | SRR9722803 | 173571 | 7425.4 |
| 136 | CC8 | 8 | t1767 | +/+ | SRR9722799 | 181743 | 6956.3 |
| 137 | CC5 | 5 | t002 | -/- | SRR9722801 | 182969 | 6684.3 |
| 138 | CC8 | 8 | t1882 | +/+ | SRR9722806 | 185318 | 7603.4 |
| 139 | CC8 | 8 | t051 | +/- | SRR9722772 | 122979 | 7258.9 |
| 140 | CC8 | 8 | t008 | +/+ | SRR9722770 | 59752 | 5687.5 |
| 141 | CC8 | 8 | t064 | -/- | SRR9722771 | 177859 | 6505.3 |
| 142 | CC5 | 5 | t002 | -/- | SRR9722767 | 171365 | 6603.3 |
| 143 | CC8 | 8 | t955 | -/- | SRR9722768 | 182810 | 7215.3 |
| 144 | CC5 | 5 | t002 | -/- | SRR9722867 | 142751 | 6883.7 |
| 145 | CC8 | 8 | t064 | -/- | SRR9722861 | 63236 | 6359.1 |
| 146 | CC5 | 5 | t002 | -/- | SRR9722859 | 154312 | 7321.4 |
| 147 | CC8 | 8 | t064 | -/- | SRR9722826 | 153034 | 6935.2 |
| 148 | CC5 | 5 | t3979 | -/- | SRR9722825 | 151850 | 6293.2 |
| 149 | CC8 | 8 | t008 | +/+ | SRR9722830 | 183027 | 6176.5 |
| 150 | CC8 | 8 | t6238 | +/+ | SRR9722828 | 171635 | 6902.7 |
| 152 | CC5 | 5 | t4371 | -/- | SRR9722893 | 149320 | 6881.0 |

| REDCap ID | Clonal Complex | MLST | Spa Type | PVL | SRR Accession | Read Number | Average read length |
|---|---|---|---|---|---|---|---|
| 154 | CC5 | 5 | t002 | -/- | SRR9722794 | 153047 | 7398.6 |
| 155 | CC8 | 8 | t051 | +/- | SRR9722896 | 130590 | 7978.0 |
| 156 | CC5 | 105 | t002 | -/- | SRR9722779 | 155036 | 7128.1 |
| 157 | CC8 | 8 | t008 | +/+ | SRR9722869 | 121934 | 7476.1 |
| 159 | CC5 | 105 | t002 | -/- | SRR9722840 | 171218 | 7946.5 |
| 160 | CC8 | 8 | t1578 | +/+ | SRR9722782 | 193963 | 7999.6 |
| 161 | CC5 | 105 | t4417 | -/- | SRR9722800 | 190963 | 7311.0 |
| 162 | CC5 | 5 | t045 | -/- | SRR9722870 | 158564 | 7959.2 |
| 163 | CC5 | 105 | t105 | -/- | SRR9722818 | 149622 | 6533.8 |
| 164 | CC8 | 8 | t064 | -/- | SRR9722823 | 141044 | 6141.6 |
| 165 | CC5 | 231 | t002 | -/- | SRR9722807 | 145069 | 6215.3 |
| 166 | CC8 | 8 | t064 | -/- | SRR9722836 | 180206 | 7002.7 |
| 168 | CC8 | 8 | t008 | +/+ | SRR9722853 | 158813 | 7154.2 |
| 169 | CC8 | 8 | t008 | +/+ | SRR9722852 | 65332 | 7486.0 |
| 170 | CC8 | 6 | t304 | -/- | SRR9722851 | 188178 | 6404.5 |
| 171 | CC5 | 5 | t002 | -/- | SRR9722850 | 171335 | 7080.5 |
| 172 | CC5 | 5 | t002 | -/- | SRR9722849 | 169657 | 7166.7 |
| 173 | CC5 | 5 | t002 | -/- | SRR9722837 | 164751 | 5985.3 |
| 175 | CC8 | 8 | t008 | +/+ | SRR9722756 | 114292 | 5923.3 |
| 176 | CC8 | 8 | t681 | +/+ | SRR9722762 | 169649 | 7134.6 |
| 179 | CC8 | 8 | t064 | -/- | SRR9722759 | 150782 | 6879.0 |
| 180 | CC5 | 105 | t002 | -/- | SRR9722763 | 150160 | 7131.9 |
| 181 | CC8 | 8 | t064 | -/- | SRR9722758 | 165583 | 7712.4 |
| 182 | CC8 | 8 | t008 | +/+ | SRR9722765 | 211386 | 7395.7 |
| 183 | CC5 | 5 | t1683 | -/- | SRR9722766 | 164963 | 7311.7 |
| 184 | CC5 | 5 | t062 | -/- | SRR9722788 | 131283 | 7848.9 |
| 185 | CC5 | 5 | t002 | -/- | SRR9722790 | 169767 | 8047.9 |
| 188 | CC8 | 8 | t211 | +/+ | SRR9722795 | 146157 | 7546.6 |
| 189 | CC8 | 8 | t068 | +/+ | SRR9722878 | 162583 | 7355.5 |
| 190 | CC5 | 5 | t002 | -/- | SRR9722881 | 102751 | 7754.3 |
| 191 | CC8 | 8 | t008 | -/- | SRR9722874 | 114151 | 7310.2 |
| 192 | CC8 | 8 | t008 | +/+ | SRR9722876 | 93558 | 7457.8 |
| 193 | CC8 | 8 | t211 | +/+ | SRR9722877 | 111155 | 7443.3 |
| 194 | CC5 | 105 | t002 | -/- | SRR9722871 | 94042 | 7291.5 |
| 195 | CC5 | 5 | t002 | -/- | SRR9722872 | 161827 | 7470.1 |
| 196 | CC5 | 105 | t002 | -/- | SRR9722901 | 177378 | 7149.9 |
| 197 | CC8 | 8 | t008 | +/+ | SRR9722900 | 172734 | 7643.1 |
| 198 | CC8 | 8 | t064 | -/- | SRR9722899 | 171666 | 6136.8 |
| 199 | CC8 | 8 | t008 | +/+ | SRR9722902 | 245476 | 4861.9 |
| 200 | CC5 | 3390 | t010 | -/- | SRR9722906 | 165922 | 6536.8 |
| 201 | CC5 | 5 | t002 | -/- | SRR9722904 | 175650 | 6305.8 |
| 202 | CC8 | 8 | t008 | +/+ | SRR9722903 | 180838 | 6415.0 |
| 203 | CC8 | 8 | t008 | -/- | SRR9722897 | 180943 | 7050.6 |
| 205 | CC8 | 8 | t008 | -/- | SRR9722847 | 194160 | 6983.3 |
| 206 | CC5 | 5 | t002 | -/- | SRR9722841 | 199323 | 7242.9 |
| 207 | CC8 | 8 | t008 | +/+ | SRR9722842 | 194193 | 7079.1 |
| 208 | CC8 | 8 | t064 | -/- | SRR9722846 | 192720 | 6960.2 |

| REDCap ID | Clonal Complex | MLST | Spa Type | PVL | SRR Accession | Read Number | Average read length |
|---|---|---|---|---|---|---|---|
| 209 | CC8 | 8 | t2658 | -/- | SRR9722844 | 195753 | 6460.6 |
| 211 | CC5 | 105 | t045 | -/- | SRR9722838 | 139873 | 7282.8 |
| 212 | CC8 | 6 | t304 | -/- | SRR9722839 | 181073 | 6965.5 |
| 213 | CC8 | 8 | t008 | +/- | SRR9722857 | 161836 | 8046.5 |
| 215 | CC8 | 8 | t008 | +/+ | SRR9722602 | 109416 | 5922.1 |
| 216 | CC5 | 105 | t4417 | -/- | SRR9722549 | 114008 | 6622.2 |
| 217 | CC5 | 5 | t548 | -/- | SRR9722572 | 196171 | 5994.2 |
| 218 | CC8 | 8 | t008 | +/+ | SRR9722594 | 178469 | 6406.4 |
| 219 | CC5 | 105 | t002 | -/- | SRR9722613 | 114050 | 5928.0 |
| 220 | CC5 | 105 | t002 | -/- | SRR9722615 | 89870 | 4497.6 |
| 221 | CC5 | 105 | t002 | -/- | SRR9722630 | 92351 | 3415.7 |
| 222 | CC5 | 5 | t002 | -/- | SRR9722633 | 147848 | 6142.5 |
| 223 | CC5 | 5 | t002 | -/- | SRR9722544 | 148150 | 6668.8 |
| 224 | CC5 | 5 | t002 | -/- | SRR9722563 | 149923 | 6999.1 |
| 225 | CC8 | 8 | t064 | -/- | SRR9722562 | 187300 | 7088.5 |
| 226 | CC8 | 8 | t064 | -/- | SRR9722656 | 165115 | 7092.6 |
| 227 | CC8 | 8 | t008 | +/+ | SRR9722655 | 155102 | 7246.5 |
| 228 | CC5 | 105 | t002 | -/- | SRR9722798 | 187625 | 7407.8 |
| 229 | CC8 | 8 | t008 | +/+ | SRR9722774 | 164956 | 6925.4 |
| 230 | CC5 | 5 | t111 | -/- | SRR9722868 | 166012 | 7496.3 |
| 231 | CC5 | 105 | t002 | -/- | SRR9722829 | 174379 | 6822.1 |
| 232 | CC5 | 5 | t062 | -/- | SRR9722791 | 134626 | 6711.3 |
| 233 | CC8 | 8 | t064 | -/- | SRR9722755 | 162021 | 7183.8 |
| 234 | CC8 | 8 | t008 | +/+ | SRR9722761 | 159240 | 6489.6 |
| 235 | CC8 | 8 | t008 | +/+ | SRR9722792 | 178569 | 7492.7 |
| 236 | CC5 | 105 | t002 | -/- | SRR9722785 | 196833 | 6952.4 |
| 237 | CC5 | 105 | t002 | -/- | SRR9722787 | 188530 | 7360.5 |
| 238 | CC5 | 5 | t002 | -/- | SRR9722854 | 156650 | 7874.9 |
| 239 | CC5 | 105 | t002 | -/- | SRR9722855 | 168459 | 7484.0 |
| 240 | CC8 | 8 | t064 | -/- | SRR9722778 | 184926 | 7638.6 |
| 241 | CC5 | 105 | t002 | -/- | SRR9722833 | 181171 | 7078.9 |
| 242 | CC8 | 8 | t008 | +/+ | SRR9722757 | 206397 | 6875.6 |
| 243 | CC5 | 105 | t002 | -/- | SRR9722821 | 203921 | 7925.4 |
| 244 | CC5 | 5 | t002 | -/- | SRR9722812 | 209665 | 8011.1 |
| 246 | CC5 | 105 | t002 | -/- | SRR9722813 | 203499 | 8203.6 |
| 247 | CC8 | 8 | t008 | -/- | SRR9722809 | 210895 | 7978.1 |
| 248 | CC8 | 8 | t008 | +/+ | SRR9722894 | 205868 | 7826.9 |
| 250 | CC5 | 5 | t002 | -/- | SRR9722773 | 205207 | 7673.7 |
| 252 | CC8 | 8 | t008 | +/+ | SRR9722775 | 189345 | 7821.1 |
| 393 | CC5 | 105 | t002 | -/- | SRR9722666 | 181783 | 7121.5 |
| 394 | CC5 | 105 | t002 | -/- | SRR9722905 | 90244 | 7796.2 |
| 395 | CC8 | 8 | t008 | +/+ | SRR9722669 | 135717 | 6577.6 |

**Supplementary Table 2. Demographic and clinical characteristics of patients with MRSA BSIs stratified by *spa* type with the odds of being USA500 vs USA300**

| | **USA500** | **USA300** | **Univariate Analysis** | | **Multivariate Analysis** | |
|---|---|---|---|---|---|---|
| **Factor** | **N = 27 (%)** | **N = 83 (%)** | **OR (95% CI)** | **p value** | **OR (95% CI)** | **p value** |
| Male | 16 (59) | 57 (69) | 0.66 (0.27-1.63) | 0.37 | | |
| *Race/Ethnicity* | | | | | | |
| Non-Hispanic White | 5 (19) | 26 (31) | Reference | | Reference | |
| Non-Hispanic Black | 11 (41) | 30 (36) | 1.91 (0.59-6.21) | 0.28 | 0.85 (0.17-4.14) | 0.84 |
| Hispanic/Latino | 9 (33) | 18 (22) | 2.60 (0.75-9.05) | 0.13 | 4.07 (0.74-22.59) | 0.11 |
| Asian | 2 (7) | 2 (2) | 5.20 (0.59-46.06) | 0.14 | 4.77 (0.39-59.03) | 0.22 |
| Unknown | 0 (0) | 7 (8) | <0.001 (<0.001->999.99) | 0.97 | <0.001 (<0.001->999.99) | 0.97 |
| *Age at Time of Infection* | | | | | | |
| 18-54 Years | 12 (44) | 38 (46) | Reference | | | |
| 55-69 Years | 11 (41) | 26 (31) | 1.34 (0.51-3.49) | 0.55 | | |
| ≥70 Years | 4 (15) | 19 (23) | 0.67 (0.19-2.35) | 0.53 | | |
| History of Injection Drug Use | 4 (15) | 12 (14) | 1.03 (0.30-3.51) | 0.96 | 0.55 (0.09-3.45) | 0.52 |
| HIV | 8 (30) | 10 (12) | **3.07 (1.07-8.85)** | **0.04** | **6.61 (1.38-31.67)** | **0.02** |
| *Admission Source* | | | | | | |
| Home | 18 (67) | 60 (72) | Reference | | | |
| NH/Rehab/LTACH | 8 (30) | 15 (18) | 1.78 (0.65-4.87) | 0.26 | | |
| Outside Hospital | 1 (4) | 8 (10) | 0.42 (0.05-3.56) | 0.42 | | |
| Prior Hospital Admission (90 Days) | 23 (85) | 47 (57) | **4.40 (1.40-13.86)** | **0.01** | **4.87 (1.15-20.55)** | **0.03** |
| *Length of Hospital Stay Prior to BSI* | | | | | | |
| ≤3 Days | 16 (59) | 58 (70) | 0.63 (0.26-1.54) | 0.31 | | |
| >3 Days | 11 (41) | 25 (30) | Reference | | | |
| *Frequent Healthcare Interaction* | | | | | | |
| Hemodialysis | 8 (30) | 14 (17) | 2.51 (0.87-7.21) | 0.09 | 1.19 (0.28-5.09) | 0.81 |
| Infusion Center | 6 (22) | 12 (14) | 2.19 (0.69-6.93) | 0.18 | 1.65 (0.38-7.17) | 0.50 |
| None | 13 (48) | 57 (69) | Reference | | Reference | |
| Presence of Invasive Device | 23 (85) | 54 (65) | 3.09 (0.97-9.79) | 0.06 | 1.50 (0.34-6.64) | 0.60 |
| Invasive Procedures | 10 (37) | 34 (41) | 0.85 (0.35-2.08) | 0.72 | | |
| Wound Present | 9 (33) | 37 (45) | 0.62 (0.25-1.54) | 0.31 | | |
| *Charlson Comorbidity Index (CCI)* | | | | | | |
| 0-3 | 5 (19) | 35 (42) | Reference | | Reference | |
| 4-5 | 11 (41) | 15 (18) | **5.13 (1.52-17.35)** | **0.009** | **9.26 (1.82-47.22)** | **0.007** |
| 6-8 | 6 (22) | 24 (29) | 1.75 (0.48-6.39) | 0.40 | 1.96 (0.43-9.05) | 0.39 |
| >8 | 5 (19) | 9 (11) | 3.89 (0.92-16.41) | 0.06 | 1.85 (0.29-11.91) | 0.52 |
| History of Transplant | 4 (15) | 10 (12) | 1.27 (0.36-4.43) | 0.71 | | |
| History of MRSA Colonization | 11 (41) | 36 (43) | 0.90 (0.37-2.17) | 0.81 | | |
| *Presumed Source of MRSA BSI* | | | | | | |
| Peripheral Intravenous Catheter | 2 (7) | 9 (11) | 0.66 (0.13-3.25) | 0.61 | | |
| Skin & Soft Tissue Infection | 3 (11) | 15 (18) | 0.57 (0.15-2.13) | 0.40 | | |
| Pneumonia | 3 (11) | 9 (11) | 1.03 (0.26-4.11) | 0.97 | | |
| Diabetic Foot Infection | 2 (7) | 6 (7) | 1.03 (0.20-5.42) | 0.96 | | |
| Vascular Access | 10 (37) | 25 (30) | 1.37 (0.55-3.39) | 0.50 | | |
| Septic Arthritis | 0 (0) | 1 (1) | -- | -- | | |
| Urinary Source | 0 (0) | 1 (1) | -- | -- | | |
| Sacral Wound | 3 (11) | 3 (4) | 3.33 (0.63-17.60) | 0.16 | **16.46 (1.98-136.87)** | **0.01** |
| Other/Unknown | 3 (11) | 9 (11) | 1.03 (0.26-4.11) | 0.97 | | |
| ICU Admission Prior to BSI | 5 (19) | 11 (13) | 1.49 (0.47-4.75) | 0.50 | | |

**Bold** = significant at ≤ 0.05
Abbreviations: NH, nursing home; rehab, rehabilitation facility; LTACH, long-term acute care hospital; BSI, bloodstream infection; HIV, human immunodeficiency virus; ICU, intensive care unit. See Table 1 for definitions.

**Supplementary Table 3. Demographics and clinical characteristics of patients with MRSA BSIs stratified by CC5/ST5 and CC5/ST105 with the odds of being CC5/ST5 vs CC5/ST105**

| Factor | CC5/ST5 N = 49 (%) | CC5/ST105 N = 61 (%) | Univariate Analysis OR (95% CI) | Univariate Analysis p value | Multivariate Analysis OR (95% CI) | Multivariate Analysis p value |
|---|---|---|---|---|---|---|
| Male | 34 (69) | 41 (67) | 1.11 (0.49-2.48) | 0.81 | | |
| *Race/Ethnicity* | | | | | | |
| Non-Hispanic White | 33 (67) | 30 (49) | Reference | | Reference | |
| Non-Hispanic Black | **2 (4)** | **18 (30)** | **0.10 (0.02-0.47)** | **0.004** | **0.09 (0.02-0.46)** | **0.004** |
| Hispanic/Latino | 11 (22) | 7 (11) | 1.43 (0.49-4.16) | 0.51 | 2.24 (0.55-9.05) | 0.26 |
| Asian | 1 (2) | 3 (5) | 0.30 (0.03-3.07) | 0.31 | 0.19 (0.01-2.62) | 0.22 |
| Unknown | 2 (4) | 3 (5) | 0.61 (0.10-3.88) | 0.60 | 0.78 (0.09-7.16) | 0.83 |
| *Age at Time of Infection* | | | | | | |
| 18-54 Years | 10 (20) | 18 (30) | Reference | | Reference | |
| 55-69 Years | 12 (24) | 18 (30) | 1.20 (0.41-3.48) | 0.74 | 0.98(0.24-4.03) | 0.98 |
| ≥70 Years | 27 (55) | 25 (41) | 1.94 (0.76-5.00) | 0.17 | 1.75 (0.34-9.08) | 0.51 |
| History of Injection Drug Use | 6 (10) | 2 (4) | 0.39 (0.08-2.03) | 0.26 | 0.26 (0.03-2.62) | 0.25 |
| HIV | 4 (7) | 0 (0) | -- | -- | | |
| *Admission Source* | | | | | | |
| Home | 20 (41) | 29 (48) | Reference | | | |
| NH/Rehab/LTACH | 19 (39) | 17 (28) | 1.62 (0.68-3.86) | 0.28 | | |
| Other Hospital | 10 (20) | 15 (25) | 0.97 (0.36-2.58) | 0.95 | | |
| Prior Hospital Admission (90 Days) | 44 (90) | 45 (74) | **3.13 (1.06-9.28)** | **0.04** | 3.36 (0.89-12.69) | 0.07 |
| *Length of Hospital Stay Prior to BSI* | | | | | | |
| ≤3 Days | 24 (49) | 30 (49) | 0.99 (0.47-2.11) | 0.98 | | |
| >3 Days | 25 (51) | 31 (51) | Reference | | | |
| *Frequent Healthcare Interaction* | | | | | | |
| Hemodialysis | 8 (16) | 10 (16) | 0.91 (0.33-2.56) | 0.86 | | |
| Infusion Center | 5 (10) | 10 (16) | 0.57 (0.18-1.82) | 0.34 | | |
| None | 36 (73) | 41 (67) | Reference | | | |
| Presence of Invasive Device | 43 (88) | 53 (87) | 1.08 (0.35-3.37) | 0.89 | | |
| Invasive Procedures | 27 (55) | 34 (56) | 0.98 (0.46-2.08) | 0.95 | | |
| Wound Present | 19 (39) | 25 (41) | 0.91 (0.42-1.97) | 0.81 | | |
| *Charlson Comorbidity Index (CCI)* | | | | | | |
| 0-3 | 11 (22) | 19 (31) | Reference | | Reference | |
| 4-5 | 10 (20) | 10 (16) | 1.73 (0.55-5.45) | 0.35 | 1.05 (0.23-4.80) | 0.95 |
| 6-8 | 20 (41) | 13 (21) | 2.66 (0.96-7.36) | 0.06 | 1.57 (0.36-6.87) | 0.55 |
| >8 | 8 (16) | 19 (31) | 0.73 (0.24-2.21) | 0.57 | 0.28 (0.05-1.72) | 0.17 |
| History of Transplant | 7 (14) | 11 (18) | 0.76 (0.27-2.13) | 0.60 | | |
| History of MRSA Colonization | 17 (35) | 26 (43) | 0.72 (0.33-1.56) | 0.40 | | |
| *Presumed Source of MRSA BSI* | | | | | | |
| Peripheral Intravenous Catheter | 3 (6) | 2 (3) | 1.92 (0.31-12.00) | 0.48 | | |
| Skin & Soft Tissue Infection | 2 (4) | 4 (7) | 0.61 (0.11-3.46) | 0.57 | | |
| Pneumonia | 2 (4) | 8 (13) | 0.28 (0.06-1.39) | 0.12 | 0.18 (0.03-1.18) | 0.07 |
| Diabetic Foot Infection | 5 (10) | 4 (7) | 1.62 (0.41-6.39) | 0.49 | | |
| Vascular Access | 14 (29) | 27 (44) | 0.50 (0.23-1.12) | 0.09 | 0.48 (0.16-1.44) | 0.19 |
| Septic Arthritis | 2 (4) | 1 (2) | 2.55 (0.23-29.02) | 0.45 | | |
| Urinary Source | 2 (4) | 1 (2) | 2.55 (0.23-29.02) | 0.45 | | |
| Sacral Wound | 5 (10) | 0 (0) | **--** | **--** | | |
| Other/Unknown | 6 (12) | 8 (13) | 0.93 (0.30-2.87) | 0.89 | | |
| ICU Admission Prior to BSI | 11 (22) | 13 (21) | 1.07 (0.43-2.65) | 0.89 | | |

**Bold** = significant at ≤ 0.05
Abbreviations: NH, nursing home; rehab, rehabilitation facility; LTACH, long-term acute care hospital; BSI, bloodstream infection; HIV, human immunodeficiency virus; ICU, intensive care unit. See Table 1 for definitions.

**Supplementary Table 4. Demographics and clinical characteristics of patients with MRSA BSIs in the HO-MRSA stratum with the odds of being CC8 vs CC5**

| Factor | CC8<br>N = 36 (%) | CC5<br>N = 59 (%) | Univariate Analysis<br>OR (95% CI) | p value | Multivariate Analysis<br>OR (95% CI) | p value |
|---|---|---|---|---|---|---|
| Male | 23 (64) | 38 (64) | 0.98 (0.41-2.32) | 0.96 | | |
| *Race/Ethnicity* | | | | | | |
| Non-Hispanic White | 10 (28) | 38 (64) | Reference | | Reference | |
| Non-Hispanic Black | 13 (36) | 10 (17) | **4.94 (1.68-14.54)** | **0.004** | **6.65 (1.68-26.36)** | **0.007** |
| Hispanic/Latino | 9 (25) | 7 (12) | **4.89 (1.46-16.36)** | **0.01** | **5.77 (1.19-27.93)** | **0.03** |
| Asian | 2 (6) | 0 (0) | >999.99 (<0.001->999.99) | 0.98 | >999.99 (<0.001->999.99) | 0.98 |
| Unknown | 2 (6) | 4 (7) | 1.90 (0.30-11.90) | 0.49 | 2.92 (0.39-22.08) | 0.30 |
| *Age at Time of Infection* | | | | | | |
| 18-54 Years | 16 (44) | 20 (34) | Reference | | | |
| 55-69 Years | 9 (25) | 15 (25) | 0.75 (0.26-2.16) | 0.59 | | |
| ≥ 70 Years | 11 (31) | 24 (41) | 0.57 (0.22-1.51) | 0.26 | | |
| History of Injection Drug Use | 3 (8) | 3 (5) | 1.70 (0.32-8.90) | 0.53 | 1.43 (0.15-13.53) | 0.76 |
| HIV | 5 (14) | 4 (7) | 2.22 (0.55-8.87) | 0.26 | | |
| *Admission Source* | | | | | | |
| Home | 24 (67) | 28 (47) | Reference | | Reference | |
| NH/Rehab/LTACH | 8 (22) | 12 (20) | 0.78 (0.28-2.22) | 0.64 | 0.88 (0.19-4.02) | 0.87 |
| Other Hospital | 4 (11) | 19 (32) | **0.25 (0.07-0.82)** | **0.02** | 0.34 (0.07-1.59) | 0.17 |
| Prior Hospital Admission (90 Days) | 26 (72) | 45 (76) | 0.81 (0.32-2.08) | 0.66 | | |
| *Frequent Healthcare Interaction* | | | | | | |
| Hemodialysis | 3 (8) | 4 (7) | 1.57 (0.32-7.67) | 0.58 | 0.49 (0.06-3.93) | 0.50 |
| Infusion Center | 12 (33) | 11 (19) | 2.29 (0.87-6.03) | 0.09 | 1.72 (0.39-7.68) | 0.48 |
| None | 21 (58) | 44 (75) | Reference | | Reference | |
| Presence of Invasive Device | 31 (86) | 54 (92) | 0.57 (0.15-2.14) | 0.41 | | |
| Invasive Procedures | 28 (78) | 44 (75) | 1.19 (0.45-3.18) | 0.72 | | |
| Wound Present | 10 (28) | 26 (44) | 0.49 (0.20-1.19) | 0.12 | 0.41 (0.11-1.51) | 0.18 |
| *Charlson Comorbidity Index (CCI)* | | | | | | |
| 0-3 | 13 (36) | 20 (34) | Reference | | | |
| 4-5 | 11 (31) | 13 (22) | 1.30 (0.45-3.77) | 0.63 | | |
| 6-8 | 9 (25) | 17 (29) | 0.81 (0.28-2.37) | 0.71 | | |
| >8 | 3 (8) | 9 (15) | 0.51 (0.12-2.26) | 0.38 | | |
| History of Transplant | 7 (19) | 12 (20) | 0.95 (0.33-2.68) | 0.92 | | |
| History of MRSA Colonization | 14 (39) | 22 (37) | 1.07 (0.46-2.51) | 0.88 | | |
| *Presumed Source of MRSA Infection* | | | | | | |
| Peripheral Intravenous Catheter | 6 (17) | 4 (7) | 2.75 (0.72-10.51) | 0.14 | **9.84 (1.46-66.50)** | **0.02** |
| Skin & Soft Tissue Infection | 4 (11) | 4 (7) | 1.72 (0.40-7.35) | 0.47 | | |
| Pneumonia | 4 (11) | 6 (10) | 1.10 (0.29-4.21) | 0.88 | | |
| Diabetic Foot Infection | 0 (0) | 2 (3) | -- | -- | | |
| Vascular Access | 19 (53) | 23 (39) | 1.75 (0.76-4.04) | 0.19 | 3.15 (0.84-11.86) | 0.09 |
| Septic Arthritis | 0 (0) | 2 (3) | -- | -- | | |
| Urinary Source | 0 (0) | 1 (2) | -- | -- | | |
| Sacral Wound | 0 (0) | 2 (3) | -- | -- | | |
| Other/Unknown | 2 (6) | 10 (17) | 0.29 (0.06-1.40) | 0.12 | 0.81 (0.09-6.96) | 0.85 |
| ICU Admission Prior to BSI | 12 (33) | 25 (42) | 0.68 (0.29-1.61) | 0.38 | | |

**Bold** = significant at ≤ 0.05
Abbreviations: NH, nursing home; rehab, rehabilitation facility; LTACH, long-term acute care hospital; BSI, bloodstream infection; HIV, human immunodeficiency virus; ICU, intensive care unit. See Table 1 for definitions.

**Supplementary Table 5. Demographics and clinical characteristics of patients with MRSA BSIs in the CO-MRSA stratum with the odds of being CC8 vs CC5**

| Factor | CC8 N = 74 (%) | CC5 N = 58 (%) | Univariate Analysis OR (95% CI) | Univariate Analysis p value | Multivariate Analysis OR (95% CI) | Multivariate Analysis p value |
|---|---|---|---|---|---|---|
| Male | 50 (68) | 40 (69) | 0.94 (0.45-1.96) | 0.86 | | |
| *Race/Ethnicity* | | | | | | |
| Non-Hispanic White | 21 (28) | 29 (50) | Reference | | Reference | |
| Non-Hispanic Black | 28 (38) | 12 (21) | **3.22 (1.34-7.76)** | **0.009** | **3.25 (1.08-9.75)** | **0.04** |
| Hispanic/Latino | 18 (24) | 12 (21) | 2.07 (0.82-5.21) | 0.12 | 2.42 (0.73-8.01) | 0.15 |
| Asian | 2 (3) | 4 (7) | 0.69 (0.12-4.13) | 0.68 | 0.64 (0.07-6.24) | 0.70 |
| Unknown | 5 (7) | 1 (2) | 6.90 (0.75-63.51) | 0.09 | 3.38 (0.27-42.76) | 0.35 |
| *Age at Time of Infection* | | | | | | |
| 18-54 Years | 34 (46) | 9 (16) | Reference | | Reference | |
| 55-69 Years | 28 (38) | 16 (36) | 0.46 (0.18-1.21) | 0.12 | 0.35 (0.10-1.31) | 0.12 |
| ≥ 70 Years | 12 (16) | 33 (57) | **0.10 (0.04-0.26)** | **<0.001** | **0.08 (0.02-0.37)** | **0.001** |
| History of Injection Drug Use | 13 (18) | 5 (9) | 2.26 (0.76-6.75) | 0.14 | 0.69 (0.16-2.97) | 0.62 |
| HIV | 13 (18) | 0 (0) | -- | -- | | |
| *Admission Source* | | | | | | |
| Home | 54 (73) | 26 (45) | Reference | | Reference | |
| NH/Rehab/LTACH | 15 (20) | 25 (43) | **0.29 (0.13-0.64)** | **0.002** | 0.50 (0.16-1.58) | 0.24 |
| Other Hospital | 5 (7) | 7 (12) | 0.34 (0.10-1.19) | 0.09 | 1.54 (0.26-8.96) | 0.63 |
| Prior Hospital Admission (90 Days) | 44 (59) | 47 (81) | **0.34 (0.15-0.77)** | **0.009** | 0.64 (0.18-2.37) | 0.51 |
| *Frequent Healthcare Interaction* | | | | | | |
| Hemodialysis | 19 (26) | 14 (24) | 1.08 (0.48-2.42) | 0.85 | | |
| Infusion Center | 6 (8) | 5 (9) | 0.96 (0.27-3.36) | 0.94 | | |
| None | 49 (66) | 39 (67) | Reference | | | |
| [a] Presence of Invasive Device | 46 (62) | 49 (84) | **0.30 (0.13-0.71)** | **0.006** | 0.64 (0.20-2.06) | 0.46 |
| [b] Invasive Procedures | 16 (22) | 21 (36) | 0.49 (0.23-1.05) | 0.07 | 0.50 (0.17-1.49) | 0.21 |
| Wound Present | 36 (49) | 22 (38) | 1.55 (0.77-3.12) | 0.22 | | |
| *Charlson Comorbidity Index (CCI)* | | | | | | |
| 0-3 | 27 (36) | 11 (19) | Reference | | Reference | |
| 4-5 | 15 (20) | 9 (16) | 0.68 (0.23-2.01) | 0.48 | 1.56 (0.35-6.95) | 0.56 |
| 6-8 | 21 (28) | 19 (33) | 0.45 (0.18-1.15) | 0.09 | 2.00 (0.50-8.00) | 0.33 |
| >8 | 11 (15) | 19 (33) | **0.24 (0.09-0.66)** | **0.006** | 1.74 (0.35-8.79) | 0.50 |
| History of Transplant | 7 (9) | 6 (10) | 0.91 (0.29-2.86) | 0.86 | | |
| History of MRSA Colonization | 33 (45) | 25 (43) | 1.06 (0.53-2.12) | 0.86 | | |
| *Presumed Source of MRSA Infection* | | | | | | |
| Peripheral Intravenous Catheter | 5 (7) | 1 (2) | 4.13 (0.47-36.37) | 0.20 | 6.13 (0.42-90.25) | 0.19 |
| Skin & Soft Tissue Infection | 14 (19) | 3 (5) | **4.28 (1.17-15.69)** | **0.03** | 2.89 (0.56-15.00) | 0.21 |
| Pneumonia | 8 (11) | 6 (10) | 1.05 (0.34-3.22) | 0.93 | | |
| Diabetic Foot Infection | 8 (11) | 7 (12) | 0.88 (0.30-2.60) | 0.82 | | |
| Vascular Access | 16 (22) | 20 (34) | 0.52 (0.24-1.14) | 0.10 | 0.78 (0.26-2.35) | 0.65 |
| Septic Arthritis | 1 (1) | 1 (2) | 0.78 (0.05-12.76) | 0.86 | | |
| Urinary Source | 1 (1) | 2 (3) | 0.38 (0.03-4.34) | 0.44 | | |
| Sacral Wound | 6 (8) | 3 (5) | 1.62 (0.39-6.76) | 0.51 | | |
| Other/Unknown | 10 (14) | 6 (10) | 1.35 (0.46-3.97) | 0.58 | | |
| ICU Admission Prior to BSI | 4 (5) | 1 (2) | 3.26 (0.35-29.96) | 0.30 | | |

**Bold** = significant at ≤ 0.05
Abbreviations: NH, nursing home; rehab, rehabilitation facility; LTACH, long-term acute care hospital; BSI, bloodstream infection; HIV, human immunodeficiency virus; ICU, intensive care unit. See Table 1 for definitions.

**Supplementary Table 6. Univariate analysis of patient outcomes**

**(A) CC8 vs CC5**

| Outcome | CC8<br>N = 110 (%) | CC5<br>N = 117 (%) | Univariate Analysis<br>OR (95% CI) | p value |
|---|---|---|---|---|
| Overall 90 Day Mortality | 23 (21) | 38 (32) | **0.55 (0.30-1.00)** | **0.05** |
| 90 Day Mortality Related to MRSA BSI | 13 (12) | 20 (17) | 0.65 (0.31-1.38) | 0.26 |
| ICU Admission Related to MRSA BSI (Missing=139) | 27 (60) | 17 (40) | 2.29 (0.98-5.39) | 0.06 |
| [a]Mechanical Ventilation Related to MRSA BSI (Missing=154) | 23 (66) | 17 (45) | 2.37 (0.92-6.10) | 0.07 |
| [b]Metastatic Infection (Missing=1) | 21 (19) | 18 (15) | 1.30 (0.65-2.59) | 0.46 |

**(B) CO-MRSA vs HO-MRSA**

| Outcome | CO-MRSA<br>N = 132 (%) | HO-MRSA<br>N = 95 (%) | Univariate Analysis<br>OR (95% CI) | p value |
|---|---|---|---|---|
| Overall 90 Day Mortality | 30 (23) | 31 (33) | 0.61 (0.34-1.10) | 0.10 |
| 90 Day Mortality Related to MRSA BSI | 20 (15) | 13 (14) | 1.13 (0.53-2.39) | 0.76 |
| ICU Admission Related to MRSA BSI (Missing=139) | 31 (79) | 13 (27) | **10.73 (3.94-29.26)** | **<0.001** |
| [a]Mechanical Ventilation Related to MRSA BSI (Missing=154) | 24 (71) | 16 (41) | **3.45 (1.30-9.15)** | **0.01** |
| [b]Metastatic Infection (Missing=1) | 31 (23) | 8 (8) | **3.34 (1.46-7.64)** | **0.004** |

**(C) CC8 vs CC5 in the HO-MRSA Stratum**

| Outcome | CC8<br>N = 36 (%) | CC5<br>N = 59 (%) | Univariate Analysis<br>OR (95% CI) | p value |
|---|---|---|---|---|
| Overall 90 Day Mortality | 11 (31) | 20 (34) | 0.86 (0.35-2.09) | 0.74 |
| 90 Day Mortality Related to MRSA BSI | 5 (14) | 8 (14) | 1.03 (0.31-3.43) | 0.96 |
| ICU Admission Related to MRSA BSI (Missing=46) | 3 (18) | 10 (31) | 0.47 (0.11-2.02) | 0.31 |
| [a]Mechanical Ventilation Related to MRSA BSI (Missing=56) | 6 (46) | 10 (38) | 1.37 (0.36-5.27) | 0.65 |
| [b]Metastatic Infection (Missing=1) | 3 (8) | 5 (8) | 0.98 (0.22-4.38) | 0.98 |

**(D) CC8 vs CC5 in the CO-MRSA Stratum**

| Outcome | CC8<br>N = 74 (%) | CC5<br>N = 58 (%) | Univariate Analysis<br>OR (95% CI) | p value |
|---|---|---|---|---|
| Overall 90 Day Mortality | 12 (16) | 18 (31) | **0.43 (0.19-0.99)** | **0.05** |
| 90 Day Mortality Related to MRSA BSI | 8 (11) | 12 (21) | 0.47 (0.18-1.23) | 0.12 |
| ICU Admission Related to MRSA BSI (Missing=93) | 24 (86) | 7 (64) | 3.43 (0.68-17.35) | 0.14 |
| [a]Mechanical Ventilation Related to MRSA BSI (Missing=98) | 17 (77) | 7 (58) | 2.43 (0.53-11.11) | 0.25 |
| [b]Metastatic Infection | 18 (24) | 13 (22) | 1.11 (0.49-2.52) | 0.80 |

Abbreviations: MRSA, methicillin-resistant *Staphylococcus aureus*; ICU, intensive care unit.
[a] Mechanical intubation excludes patients who were perioperative that are intubated < 4 days.
[b] Metastatic infection is defined as evidence of bacterial seeding to other body sites after initial bloodstream infection (i.e. endocarditis, spinal infection, septic pulmonary emboli).

**Supplementary Figure 1. Gene distribution by patient zip code.** Geographic information system (GIS) map of New York City illustrating the distribution of each patient isolate by CC. The map excludes 40 isolates that were located outside the New York City borough boundary. Of the 185 isolates mapped, 96 belong to CC5 and 89 belong to CC8.

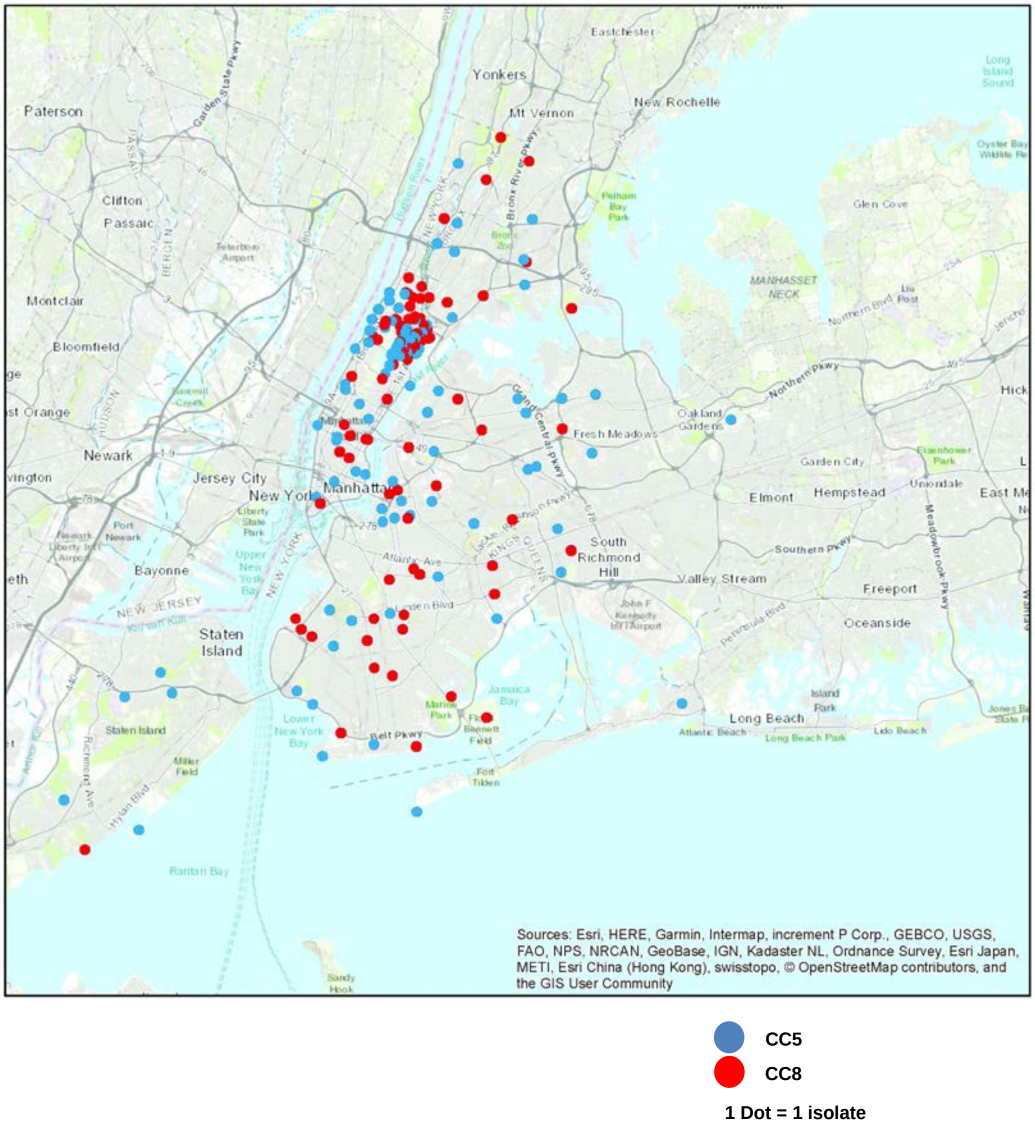

**Supplementary Figure 2. Death related to MRSA BSI within 90 days stratified by clone.** Survival analyses of CC8 and CC5 90-days from the first positive MRSA culture. Tick marks represent censored patient due to discharge from the hospital. Risk tables reflect the number of patients still at risk of death at 30, 60, and 90 days.

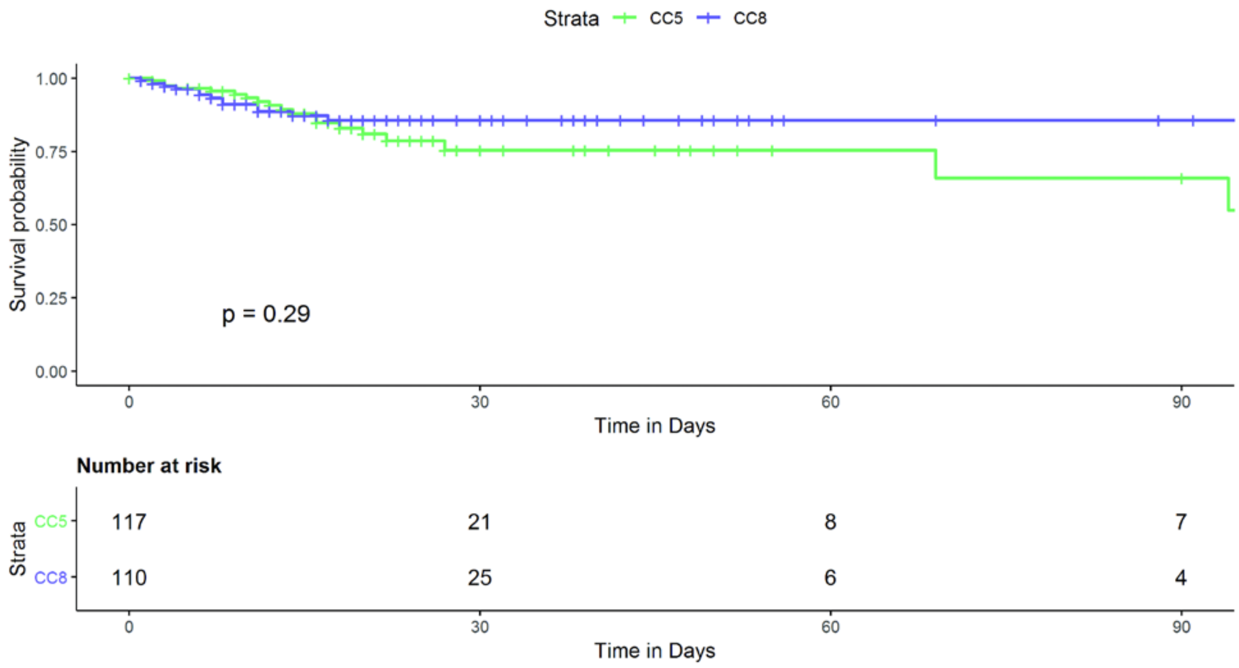

**Supplementary Figure 3. Multivariate analysis of death related to MRSA BSI within 90 days.** Forest plot of the multivariate Cox regression for all variables that had *p*≤0.2 in the univariate analysis. The square represents the hazard ratio (HR), and the lines reflect the 95% confidence interval.

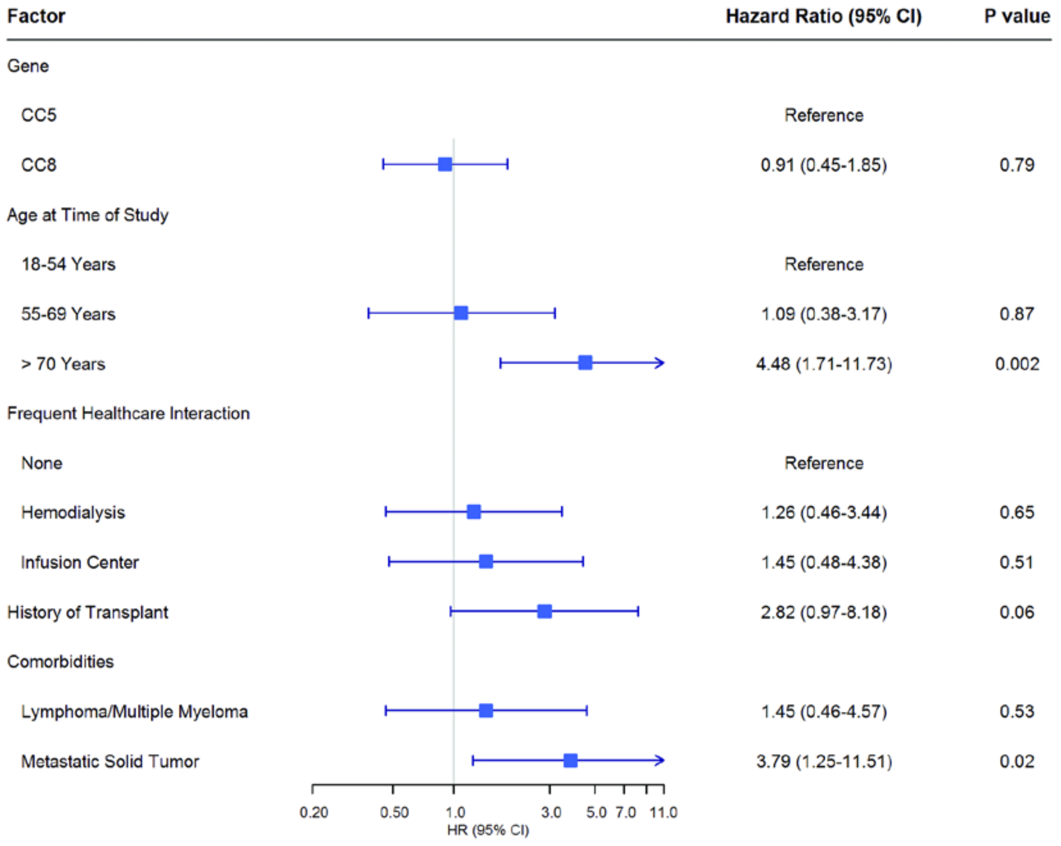
